# Microbial community organization during anaerobic pulp and paper mill wastewater treatment

**DOI:** 10.1101/2023.08.11.553022

**Authors:** Torsten Meyer, Minqing Ivy Yang, Camilla Nesbø, Emma Master, Elizabeth A. Edwards

## Abstract

Amplicon sequencing data and operating data from anaerobic wastewater treatment plants from three Canadian pulp and paper mills were explored using correlation and network modularization approaches to study the microbial community organization and identify relationships between organisms and operating conditions.

Each of the digesters contain two or three modules consisting of organisms that cover all trophic stages of anaerobic digestion. These modules are functioning independently from each other, and their relative abundance changes in response to varying operating conditions.

The time delay between a change in digester operation and the change in the abundance of microorganisms was investigated using time-lagged operating parameters. This time delay ranged between two to four days and is likely influenced by the growth rates of the anaerobic microorganisms and the digester hydraulic retention time.

Digester upsets due to plant shutdown periods and organic overload caused a drastic increase in the population of acetoclastic methanogens, acidogenic fermenters, and syntrophic acid degraders. As a response to impaired process conditions, the same *Methanothrix* amplicon sequence variant (ASV) dominated methanogenesis in the digesters of all three mills. The common characteristics of the organisms represented by this ASV should be further investigated for their role in alleviating the impact of digester upset conditions.

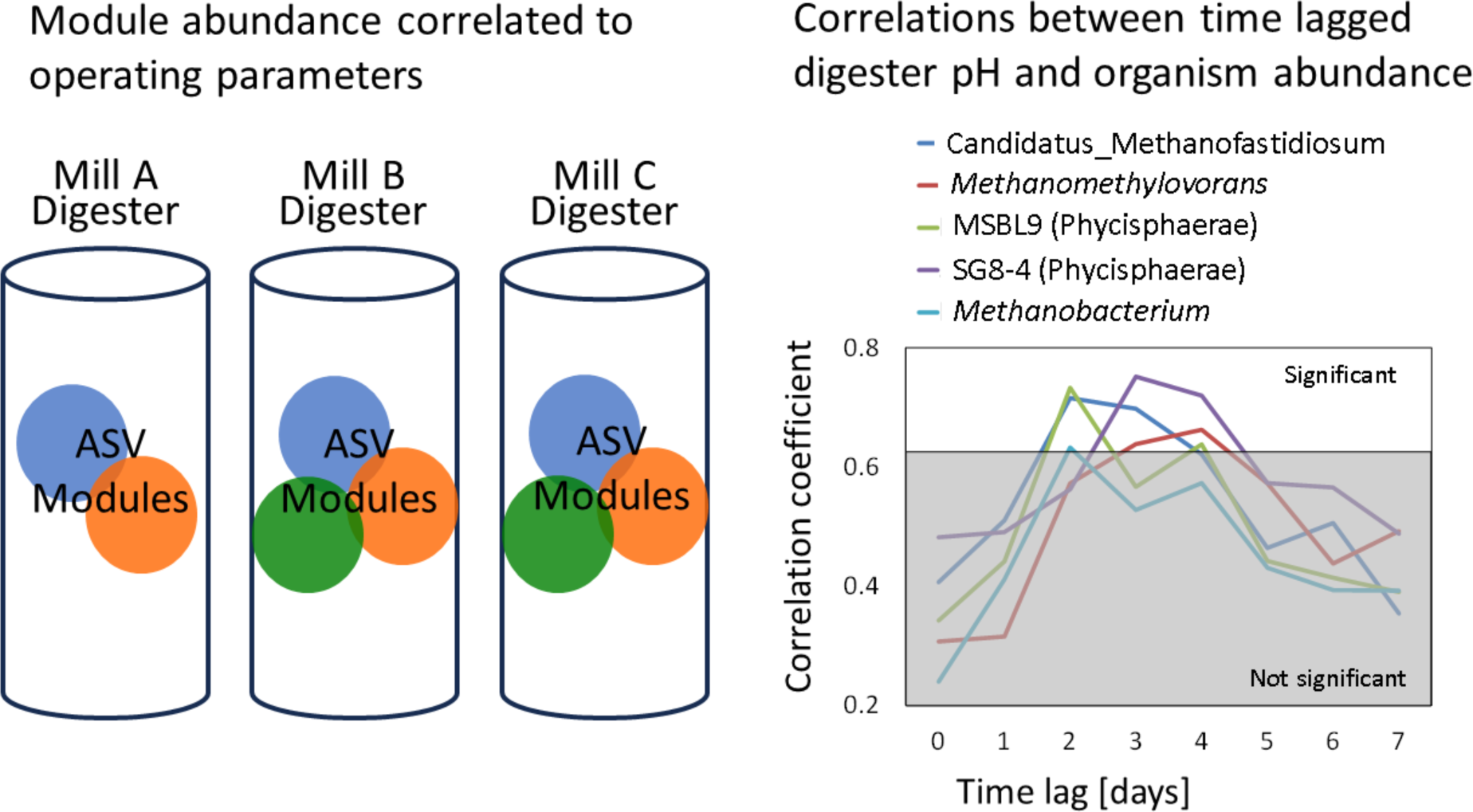

## 1 Introduction

Anaerobic digestion has been widely applied for the treatment of high-strength industrial wastewater. However, digesters are still often experiencing phases of process deterioration where the root causes are not completely clear. The process impairment may be related to the presence of toxicants or inhibitors in the wastewater, organic overload, the lack of essential nutrients,^1^ or large fluctuations in the composition of the wastewater. The roles that the microbial community plays have for a long time been the last frontier of knowledge about anaerobic digestion. As sequencing methods have improved and become more affordable much insight has been gained in the interactions between microbial community and digester operation. Numerous studies have investigated community dynamics during anaerobic digestion of organic solids and wastewater. Most of this research is based either on bench-scale experiments where operating parameters were systematically changed to investigate the response of the microbial community, or full-scale studies where microbial communities were observed and related to the varying operating conditions.^2–10^ These studies have significantly enhanced our understanding of anaerobic digestion however, there are still large uncertainties, and the results of previous studies are to some extent contradictory. Dennehy et al.^2^ found that despite of changes of operational parameters such as hydraulic retention time and the ratio between the feed substrates pig manure and food waste the microbial profile did not change significantly. On the other hand, De Vrieze et al.^9^ observed constant evolution of the microbial communities over time in several full-scale digesters where the impact of the operational conditions on these dynamics is relatively small. Wu et al.^10^ also observed ongoing changes of the microbial community although the digester operating conditions and process performance were kept constant and stable.

Few studies have explored modules of microorganisms within a digester that have meaningful biological functions. The identification of microorganism grouping, in form of modules, can enhance our understanding of the interplay among these organisms during anaerobic wastewater treatment. Modules may be described as groups of closely linked microorganisms that act as functional units and interact with each other within an ecosystem. Wang et al.^11^ detected two modules in water-submerged paddy soil samples, one associated with aerobic respiration and fermentation, and the other with metal/sulfur cycles. Similarly, in a study on anaerobic digestion of antibiotics, Zhang et al.^12^ identified two modules within a digester consisting of Firmicutes and Bacteroidetes that were positively correlated with antibiotics removal. In our research, we have identified distinct modules within each digester that function relatively independently and show correlations with several operating parameters.

Calculating correlations and co-occurrences between microorganisms and operating parameters is one way with which functional relationships in anaerobic digesters may be identified. However, revealing meaningful relationships within a microbial community is difficult for a variety of reasons. Uneven sequencing depth, the occurrence of rare organisms with a high fraction of zero counts, and compositionality are serious challenges and can lead to false conclusions.^13^ Compositionality is associated with an inherent negative correlation structure when community datasets are analyzed based on relative abundance. Because the abundance of each ASV is divided by the total abundance of all bacterial or archaeal ASVs in a sample, an increase in abundance of one ASV must decrease the abundance of others. In many cases amplicon sequencing data consist of time series and identifying correlations between time series is associated with additional challenges. Time series characteristics such as autocorrelation, non-stationarity, and seasonality can generate spurious correlations or inflate existing correlations when applying standard statistical methods.^14^

Microbial communities in anaerobic digesters exhibit large numbers of potential relationships. Therefore, computational methods to generate correlation networks have been increasingly applied. Because of the difficulty and uncertainty in identifying meaningful microbial relationships, Weiss et al.^13^ compared the performance of the most widely used correlation network techniques, including Correlation Networks (CoNet), Local Similarity Analysis (LSA), Maximal Information Coefficient (MIC), Random Matrix Theory (RMT), Sparse Correlations for Compositional data (SparCC), and the naïve correlation coefficients Bray-Curtis, Pearson, and Spearman. Performance comparison was implemented based on a large number of different artificial and real datasets reflecting a wide variety of environmental scenarios. Of all investigated techniques, LSA appeared to be the most suitable method for identifying correlations between time series such as the relative abundance of ASVs over time. LSA is unique in that it can identify local and time-delayed correlation and association structures in microbial community datasets.^15^ The “local” in LSA refers to associations between two organisms that may occur only for part of the time period of investigation. Also, LSA has shown to be able to identify competitive three-species relationships which involve both co-existence and mutual exclusion.^13^ These relationships have been previously interpreted as correlations between two organisms that are mediated by a third organism or an environmental parameter.^16^ Although three-species relationships can be difficult to interpret, they may enhance the understanding of the synergistic or competitive dynamics that may influence the efficacy of anaerobic treatment processes.

To identify time-delayed associations between environmental parameters, cross-correlation analyses are commonly applied. The time lag with the highest correlation coefficient is usually considered as the true response time between two time series (Olden et al.,^17^ and refs therein). However, autocorrelation may cause discrepancies between the true time lag and the calculated lag, and/or inflate the number of false positive findings, i.e. for microbial community data, the erroneous identification of associations between two organisms. Another potential source of error is based on intra-multiplicity, which refers to the identification of spurious cross-correlations by chance, due to a large number of examined time lags between two time series. These errors are usually smaller with a decreasing autocorrelation, a decreasing number of applied time lags, and an increasing true correlation between the two time series.^17^ Because of the cost and labour required for amplicon sequencing, most previous studies have applied relatively sparse datasets at intervals of weeks, months or years. However, the response time between changes in environmental parameters and microorganism growth is usually much shorter and in the range of hours to several days. Therefore, in contrast to previous studies we have used smaller intervals by generating time lagged versions of daily measured operating parameters and have correlated these to the sequencing data.

The first section of the paper discusses the community profile and the most important organisms in each of the digesters. The second section is dedicated to the description of large and independently functioning modules of ASVs within the reactors. Three-species relationships involving individual organisms and operating parameters are discussed in the third section. Results from the correlation analysis using lagged parameters are presented in the fourth section. Finally, the fifth section discusses the response of key organisms to digester upset events.

## 2 Experimental/Methods

### 2.1 Mill operating data

Mill A combines a chemical pulp mill and a bleached chemical thermo-mechanical pulp (BCTMP) mill, and mills B and C are both BCTMP mills. The three mills operate different types of anaerobic treatment reactors. Mill A operates two internal circulation (IC) reactors in parallel with a hydraulic retention time (HRT) of 8 – 12 hours, fed with three streams consisting of BCTMP wastewater, acid condensate, and an alkaline filtrate, all of which have highly varying flow rates and composition (Figure 1). Mill B operates three anaerobic hybrid digesters in parallel with an HRT of 2 – 3 days, fed with composite BCTMP wastewater. Mill C operates an anaerobic lagoon with an HRT of 12 – 14 days, which is also fed with composite BCTMP wastewater. Daily operating data from these reactors from January 2017 to August 2018 were compiled. The data consist of numerous parameters from each mill including soluble/total chemical oxygen demand (COD) removal efficiency, volatile fatty acid (VFA) to alkalinity ratio, hydraulic retention time, organic loading rate, wastewater flow rates, influent/effluent concentrations of total suspended solids, sulfur compounds, COD, and others (Tables S13-S15). Some known inhibitors to anaerobic digestion of pulp mill wastewater are undissociated sulfide and sulfite as well as resin and fatty acids (RFAs). The sulfur compounds are measured differently in the three mills. Mill A analyzes sulfate and sulfite concentrations in the influent on a daily basis, mill B measures total sulfur levels in the influent and H_2_S concentrations in the biogas daily, and mill C measures sulfate concentrations on a weekly basis, and H_2_S concentrations in the biogas approximately once every other day. RFAs are analyzed occasionally in all mills.

**Figure 1.**
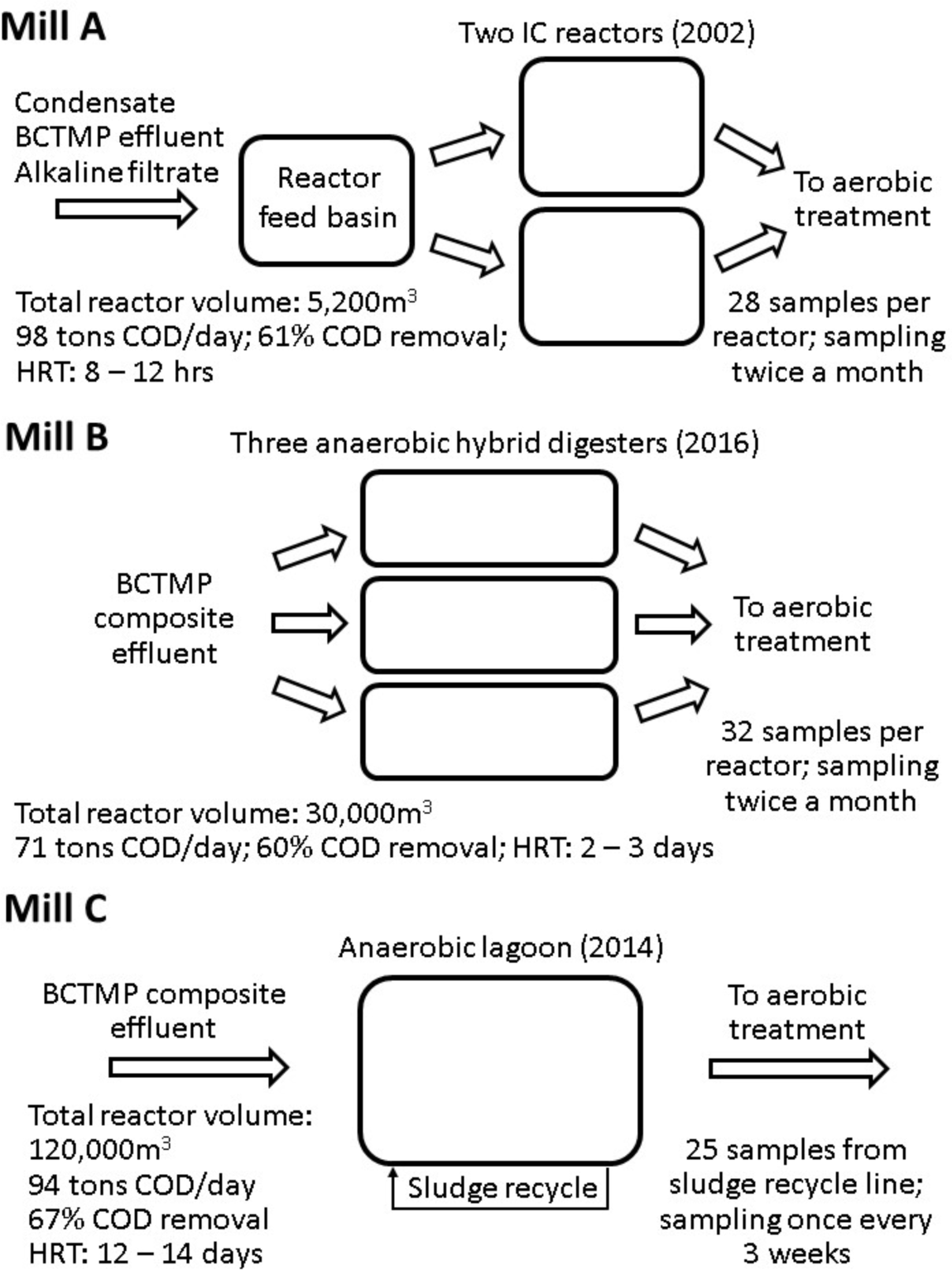
Schematic of the anaerobic reactors in the three mills, average operating parameters, and the years of reactor commissioning.

### 2.2 Sampling and DNA extraction

Samples of anaerobic biomass were collected approximately twice a month over the course of 1.5 years from the anaerobic treatment reactors at the three mills. The samples from mill A were collected from a sampling port 6 m above the bottom of the IC reactor, the samples from mill B were taken from a sampling port 0.5 m above the reactor bottom, and the samples from mill C were taken from the anaerobic return sludge line. The samples from mill A were frozen and shipped whereas the samples from mills B and C were cooled and shipped to University of Toronto. Upon arrival the samples from mills A (after thawing), B and C were centrifuged, and the pellets were stored in the freezer at −80°C until DNA extraction. The number of collected samples is 56 (28 per digester) from mill A, 96 (32 per digester) from mill B, and 25 from mill C. Total community DNA was extracted using the Power Soil DNA Isolation Kit (MoBio Laboratories, Carlsbad, CA) from 10 g of sample. The quantity and quality of DNA extracts were confirmed using NanoDrop spectrophotometer ND-1000 (Thermo Fisher Scientific, Wilmington, DE). All DNA extracts were stored at –80◦C.

### 2.3 Microbial community analysis

DNA extracts were sent to the McGill University and Genome Quebec Innovation Center (Quebec, Canada) for amplicon sequencing using the Illumina MiSeq system and the V3 reagent kit with primers targeting the V6-V8 regions of the 16S rRNA gene, with sequences 926f-modified: 5’-AAACTYAAAKGAATWGRCGG-3’ and 1392r-modified: 5’-ACGGGCGGTGWGTRC-3’.^18^ The raw amplicon sequence data were submitted to the National Centre for Biotechnology Information’s sequence read archive database under BioProject PRJNA916529.

The raw amplicon sequences were processed and analyzed using QIIME2 version 2019.10.^19^ After trimming the primer region with the cutadapt plug-in, amplicon sequence variants (ASVs) were generated using the DADA2 plug-in with the following settings: p-trunc-len-f = 260, p-trunc-len-r = 240 or 220 bp, p-max-ee = 2. The amplicon sequencing read numbers for each QIIME step are provided in Table S12. The resulting data set was subsampled to an equal depth of 28,117 reads per sample prior to analysis to minimize the bias caused by different read-depth. Taxonomic classification was performed using the Silva-132-99-nb classifier trained on the 926f and 1392r primer set. All ASVs are assigned a feature number at the end of the name to distinguish between different ASVs with the same taxon name. The relative abundance data for all digesters are provided in excel Tables S5-S10, and representative sequences are in Table S11. Due to strong correlations observed in the operating and ASV data among the in parallel operating digesters, the results presented in sections 3.2 - 3.5 represent only one digester from mill A and mill B.

### 2.4 NMDS and heatmaps

Non-metric multidimensional scaling (NMDS) plots were generated to condense the multi-dimensional data into a two-dimensional representation using Ampvis2 in R.^20^

Heatmaps based on relative abundance data were generated for the twenty most abundant archaea and bacteria in each mill. In the case of mill A and mill B, the maps show the average abundance in all digesters within one mill. The heatmaps were generated using the seaborn module in python.^21^

### 2.5 Correlation analysis and module identification

The analytic software eLSA (Extended Local Similarity Analysis)^15^ was used for the correlation calculations and is available for free download (http://bitbucket.org/charade/elsa). The applied LSA pipeline includes f-transformation, normal score transformation, correlation calculation, and statistical significance evaluation. Local similarity (LS) scores between two series, i.e. two ASVs, were calculated by means of Pearson correlation, following normalization.^22^ P-values were determined through permutation, and by the proportion of correlation scores larger than the original score after shuffling one of the two series 1000 times and re-calculating the score each time. Multiple hypothesis correction was done with false discovery rates, or q-values.^23^ The software also enables the calculation of Liquid Associations (LA) which can identify three-feature relationships. The term liquid (as opposed to solid) refers to the correlation strength between two organisms (X, Y) that is mediated by a third organism or an environmental variable (Z). Assuming that X, Y and Z are three standard normal distributed variables with a mean 0 and variance 1, the liquid association score of X and Y conditionally on Z is denoted by LA(X,Y | Z). Mathematical derivations by Stein^24^ and Li^22^ lead to LA(X,Y | Z) = E(X,Y,Z), which is the mean (or expected value) of the product of the three normalized series. Accordingly, LA scores were calculated in two steps. First, the data were standardized by means of normal score transformation, and second the average product of X, Y, Z was calculated. The variable Z can mediate correlations between the variables X and Y in four different scenarios. A high level of Z may increase the positive correlation between X and Y or increase the negative correlation between X and Y. Likewise, a low level of Z may increase the positive correlation between X and Y or increase the negative correlation between X and Y^16^. The relative abundance based on the overall bacterial and archaeal community in each sample was used for all correlation calculations.

The cytoscape bioinformatics platform and the plug-in ModuLand^25–27^ were used to identify modules consisting of organisms that are related to each other. ModuLand determines modular assignment values to each of the organisms within a module depending on the degree of “belonging” to that module (Tables S2-S4). The latter is based on local similarity (LS) scores that were determined by LSA. The modules overlap with each other, i.e. many of the organisms were assigned to more than one module. However, the highest modular-assignment-values determine the modules to which an organism was assigned. Of the modules identified by the algorithm only those that contained at least ten organisms with modular assignment values greater than 4.0 were selected because these modules represent relatively independently functioning clusters of ASVs. In most cases, the organisms in the selected modules could be unequivocally assigned to only one module or very few overlaps with other modules were identified. Also, only organisms with a relative abundance >1% in at least one sample were included. All modules were subsequently analyzed in terms of associations with digester operating parameters (Tables S13-S15) to explore the functional relationships between individual modules and the anaerobic treatment conditions. Because operating parameters often exhibit non-linear distributions, spearman correlations were applied as correlation measure for these relationships. Also, because of the time lag between changes in operating conditions and ASV abundance (see sections 2.6, 3.4), ASV abundances were correlated with operating parameters lagged by 2 days (Figure S7). A relationship between a module and a lagged operational parameter (see section 2.6) was established when 100% of all significant correlations (p<0.05) between the operating parameter and the ASVs within a module were either positive or negative, or in the case of module B1, >90% of all significant correlations were either positive or negative (Tables S16-S18 in Supporting Tables).

### 2.6 Lagged parameters

To investigate the delayed response of microorganism abundance to changes of wastewater treatment parameters, lagged operating parameters were generated. Time series of daily operating parameters were shifted forward in time by one to seven days and added to the existing dataset, i.e., variables were added at day t + i for i = 1 to 7 days (Figure S7). The variable i refers to the response time of the ASV abundance as a result of changes in operating parameters. Thereby, the relative abundance of microorganisms on the day of sampling could be correlated to operating data that were measured 1 to 7 days ago.

In addition to computing LS scores, LSA also calculates Spearman correlations, with p-values determined through permutation. To identify the true time lag between operational changes and changes in microbial abundance, as well as minimize false discovery rates of the lagged parameter associations, only highly significant Spearman correlations with p<0.001 and q<0.05 were used. Also, operating parameters were lagged by only seven days.

### 2.7 Digester upsets

Periods of digester performance impairment were assessed to investigate the response of individual ASVs to adverse conditions. These periods were identified by means of a combination a high VFA-alkalinity ratio and low COD removal efficiency. The VFA-alkalinity ratio is a digester control parameter where elevated values may indicate imminent process instability. Within the timeframe of the study (Jan 2017 – Aug 2028), only one major upset was identified in each of the mills A and C, which were considered for this study. While mill B experienced multiple smaller upsets, there was only one event characterized by a COD removal efficiency drop below 35% and an increase in VFA-alkalinity ratio above 0.5, and this upset period was used for the investigation.

## 3 Results and Discussion

### 3.1 NMDS, heatmaps, and most abundant ASVs

NMDS plots were generated for archaea and bacteria from the digesters of the three mills (Figure 2). The points representing the samples from both reactors at mill A are very close to each other indicating that the microbial community is similar in both digesters and did not change much during the time period of investigation. On the other hand, the points related to the samples from mill B are notably further apart from each other, particularly for the archaeal community, highlighting the relatively large dissimilarity between the samples. The sample dissimilarity at mill C is larger than at mill A but smaller than at mill B. These results may be explained by the different degree of acclimation and adaption of the microbial communities in the digesters. Prior to this study, the anaerobic reactors at mill A were in operation for 15 years, and the community was likely well adapted to the mill wastewater. In contrast, the digesters at mill B were put into operation three months before the sampling campaign started, and the microbial community has not had much time to adapt to the substrate contained in the mill wastewater. The anaerobic lagoon at mill C was in operation for three years prior to the onset of this study. It seems that a longer adaptation time leads to a more stable microbial community. According to anecdotal evidence from personnel at mill A, after the reactors were commissioned the adaptation process has been progressing over the course of several years, leading to an increasing resistance of the microbial community to anaerobic inhibitors. The similarity between the samples from both reactors at mill A indicates that the two communities respond very similar to the varying wastewater composition, possibly because they are operated in the same way.

**Figure 2.**
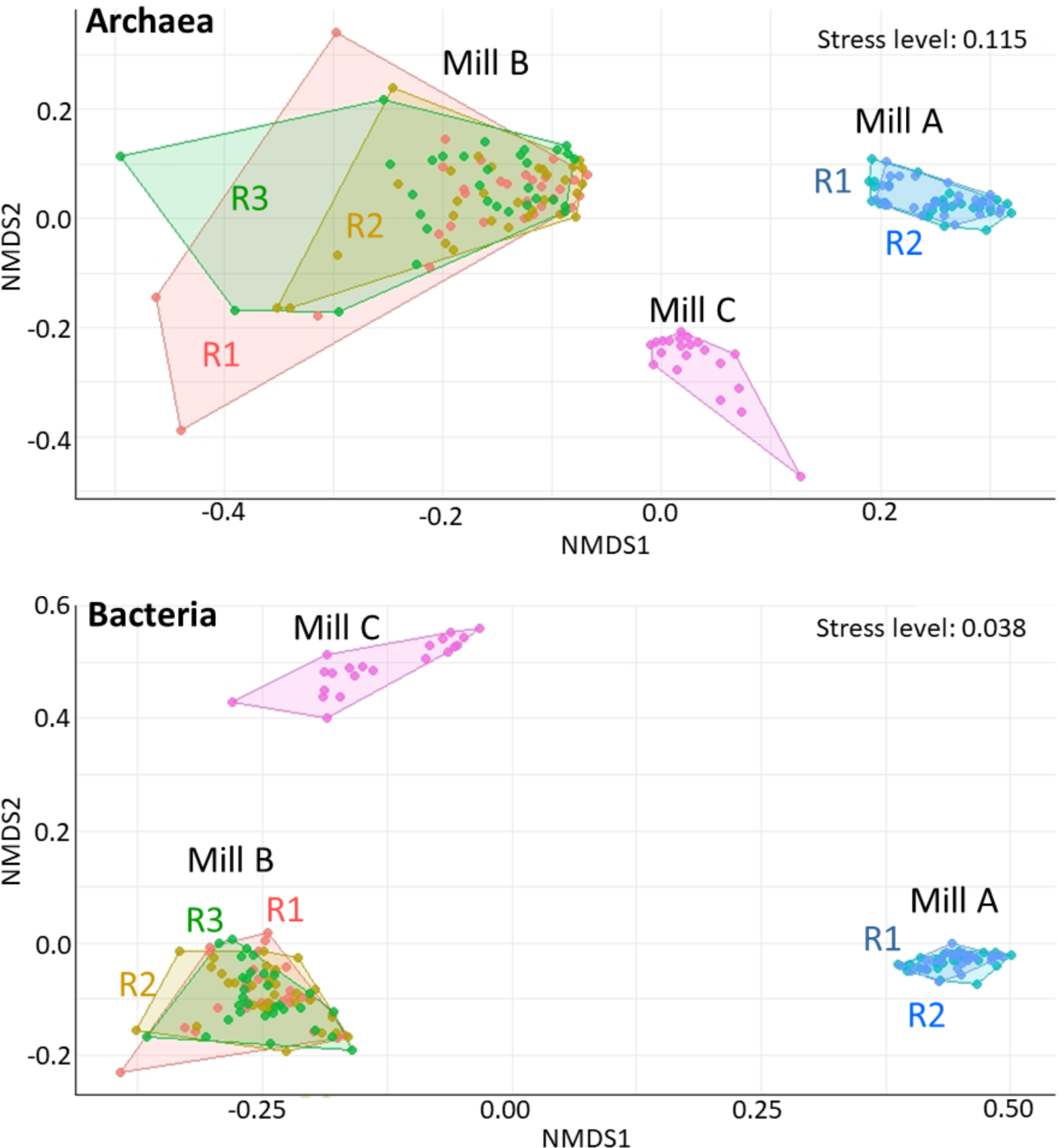
Non-metric multidimensional scaling (NMDS) ordination of ASV data from the digesters in the three mills. Mills A and B comprise multiple reactors (R).

Heatmaps including the twenty most abundant archaeal and bacterial organisms in the digesters are shown in Figures 3 for Mill A, S1 for Mill B, and S2 for Mill C. The heat maps for mills A and B show abundances averaged across the 2 or 3 digesters within each mill. Conversely, all other abundance values presented in this paper exclusively belong to only one digester. At mill B the abundance of the most common ASVs vary to a larger extent than at the other two mills. This corroborates the results from the NMDS analysis indicating a higher degree of microbial community stability in the reactors at mills A and C.

**Figure 3.**
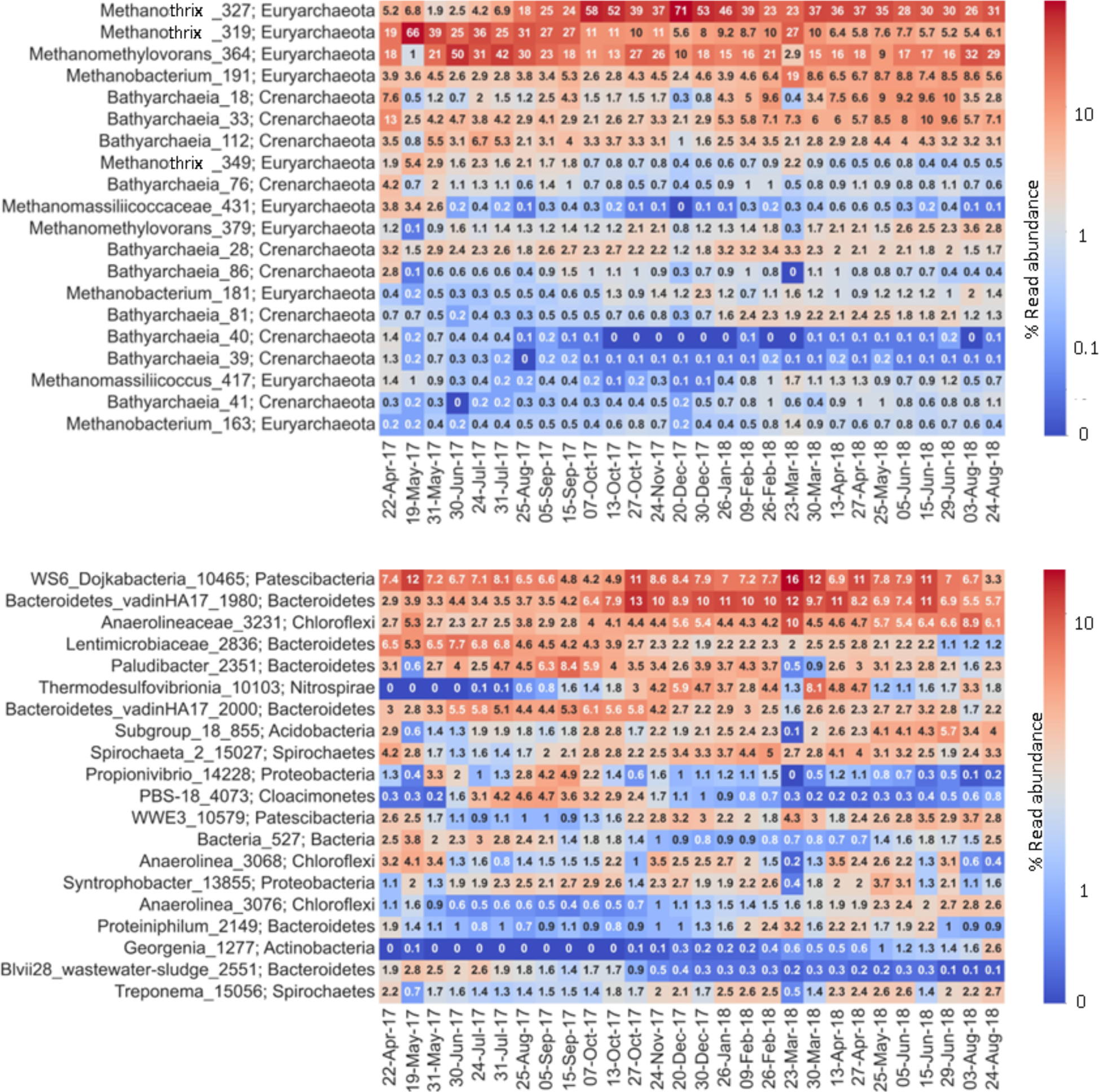
Heatmaps of the twenty most abundant archaea (top) and bacteria (bottom) in the anaerobic reactors of mill A (based on the average relative abundance in both digesters). Similar heatmaps for mills B and C are shown in Figures S1 and S2

ASVs with an abundance of >5% in at least one sample were considered high-abundance organisms and are discussed in the following. Acetoclastic methanogenesis dominates biogas production in all three mills, and ASVs of *Methanothrix* are, on average the most abundant methanogenic species (average over the period of investigation: mill A - 63%, mill B - 52%, mill C - 73%) (Figures 3, S1, S2). The reactors also contain high-abundance hydrogenotrophic and methylotrophic methanogens, including ASVs of *Methanomethylovorans* and *Methanobacterium* at mill A, *Methanomethylophilaceae*, *Methanomethylovorans* and *Methanofollis* at mill B, and *Methanomethylovorans* and *Methanospirillum* at mill C. Also, each reactor contains at least three ASVs of high abundance from *Bathyarchaeia*. In all digesters the fraction of the total archaeal community consisting of *Bathyarchaeia* ranges between 10% and 20%. Members of this class have been associated with a variety of functions including methane metabolism, cellulose hydrolysis and fermentation, and reductive acetogenesis.^28–31^ These non-methanogenic archaea have also been associated with lignin degradation^32,33^ which would explain their high relative abundance in the mill digesters. Pulp mill wastewater contains high concentrations of lignin derived compounds as a result of the pulping process and the associated separation of lignin from wood. In this study, *Bathyarchaeia* are significantly correlated to hydrogenotrophic and methylotrophic methanogens (Table S1). *Bathyarchaeia* may produce hydrogen to be consumed by these methanogens, or they prefer similar conditions as these methanogens by utilizing hydrogen and/or methylated compounds. Yu et al.^33^ suggest *Bathyarchaeia*-mediated lignin demethylation as a key step in lignin degradation.

The most abundant bacterial ASVs in all mills are classified to the phylum Bacteroidetes, however, Spirochaetes, Chloroflexi, Proteobacteria are also present at notable abundance. At mill A, high-abundance bacteria are ASVs of *Ca. Dojkabacteria, Bacteroidetes_vadinHA17, Anaerolineaceae, Lentimicrobiaceae, Paludibacter, Thermodesulfovibrionia, Subgroup_18 (*phylum Acidobacteria), and *Spirochaeta*. The majority of these bacteria are hydrolyzers and fermenters/acidogens indicating that the wastewater contains complex organic material that these organisms break down into smaller compounds such as acids and other fermentation products.

The two bacteria with the highest average abundance, *WS6_Dojkabacteria_*10465 from the phylum Patescibacteria and *Bacteroidetes_vadinHA17_*1980 have been previously associated with hydrolysis^34,35^. The most abundant fermenters include ASVs of *Lentimicrobiaceae*, *Anaerolineaceae* and *Paludibacter*. Members of the family *Lentimicrobiaceae* degrade carbohydrates and produce volatile fatty acids (VFA).^36^ *Paludibacter* use sugars to produce acetate and propionate.^37^ Members of the Anaerolineae class have been referred to as one of the core populations in anaerobic digesters. Some of them have been associated with cellulose degradation and anaerobic syntrophy^38^ with hydrogenotrophic methanogens.^39^ Due to its filamentous morphology, *Anaerolineae* organisms may play a role in the formation of anaerobic granules in high-rate UASB-type reactors^40^ such as the IC reactors at mill A.

At mill B, high-abundance bacterial ASVs are classified as *Proteiniphilum, Dysgonomonadaceae, Bacteroides, Paludibacter, Sphaerochaeta, Anaerolineaceae, Prevotellaceae, Prevotella*, and *Ruminococcaceae*. All these organisms, except the ASVs from the family *Prevoteallaceae* and *Ruminococcaceae*, have been associated with fermentation or acidogenesis^38, 41–45^. The predominance of fermenting bacteria may be an indication that the organic matter in the digester feed at mill B is somewhat less complex and of smaller molecular size than at mill A. The alkaline filtrate at mill A, as part of the wastewater, contains elevated levels of high-molecular lignin-derived compounds and polysaccharides.

High abundance bacteria at mill C are *Paludibacter, Prolixibacteraceae, Smithella, Syntrophus, Sphaerochaeta, Bacteroidetes_vadinHA17, Subgroup_18 (*phylum Acidobacteria*), Ercella,* and *LD1-PB3 (Verrucomicrobiae).* The fermenter *Paludibacter*_2351 ^44^ and the hydrolyzer *Prolixibacteraceae_*2527 ^46^ dominate the bacterial community. Both *Smithella* and *Syntrophus* have been associated with acetogenesis,^47–49^ and Bacteroidetes_vadinHA17 has hydrolyzing capabilities.^35^ *Ercella* and *LD1*-*PB3* (Verrucomicrobiae) may be involved in more than one anaerobic digestion stage. Both have been previously associated with hydrolysis and fermentation/acidogenesis, the latter also with acetogenesis.^5,50^ The digester community at mill C contains high-abundance ASVs from all stages of anaerobic digestion.

### 3.2 Biologically meaningful modules identified based on Local Similarity Analysis

Modules consisting of organisms that have covarying relative abundances were identified based on local similarity (LS) scores. Large, stable, and relatively independent modules consisting of organisms with >1% relative abundance in at least one sample were identified. When applying the ModuLand algorithm to these large and diverse datasets, relatively few clusters or modules of organisms emerged for each mill. Each of these modules grouped phyla that span the trophic stages of anaerobic digestion. The modules correlated well with specific mill operating parameters. Each module also comprised phyla at relatively high abundance, and the modules within a mill were independent of each other, further supporting their meaningful association to biological functions.

#### 3.2.1 Modules in mill A

The analysis of the digester community in mill A returned two modules that fulfill the conditions described in the methods section (Figure 4). Each of these modules contained hydrolyzers, fermenters/acidogens, acetogens, and methanogens, and therefore covered all stages of the anaerobic digestion process.

**Figure 4.**
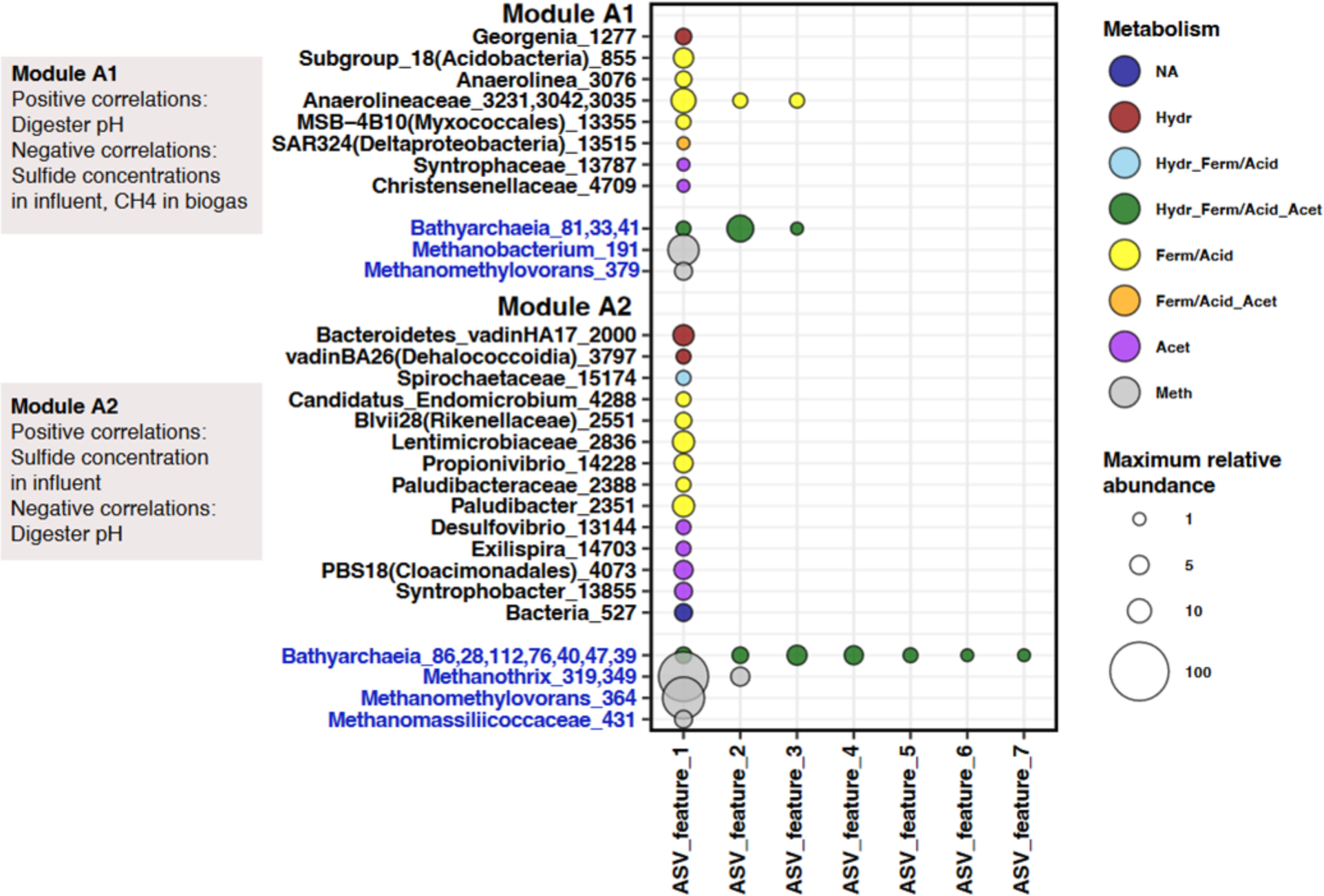
Modules in digester 1 at mill A and correlations with operating parameters. For each module ASVs with the same taxonomic classification are shown on the same line. Black names are bacteria; blue are archaea. Numbers after taxonomy label refers to the ASV feature id in Tables S5-10. Abundances of up to 7 ASVs with the same taxonomy are shown as circles where size refers to relative abundance of the ASV (bacteria and archaea normalized separately). Hydr – Hydrolysis, Ferm – Fermentation, Acid – Acidogenesis, Acet – Acetogenesis, Meth – Methanogenesis.

The two modules are correlated to several operating parameters. The strongest correlations appeared to be with the oxidation-reduction potential (ORP) in the reactor feed basin. This parameter is negatively correlated to the sulfide content in the digester feed, i.e., a low ORP means high sulfide concentrations in the influent. Module A1 is positively correlated to the digester influent pH, and negatively correlated to the sulfide concentrations in the influent and the methane content in the biogas. On the other hand, module A2 is positively correlated to the sulfide content in the influent, and negatively correlated to the pH (Table S16). This suggests that module A2 is predominant under environmental stress conditions caused by high sulfide concentrations. On the contrary, module A1 seems to prevail when the treatment conditions are more stable. Several high abundance organisms are not part of these two modules, including *Methanothrix_*327 (73%), *Bathyarchaea_*18 (14%), *WS6_Dojkabacteria_*10465 (16%), *Bacteroidetes_vadinHA17_*1980 (14%*), Thermodesulfovibrionia_*10103 *(8%),* and *Spirochaeta_*15027 (5%). The corresponding organisms are likely more flexible in terms of their dependence on other community members and/or environmental conditions.

#### 3.2.2 Modules in mill B

Three modules were identified as part of the microbial community at mill B (Figure 5). Similar to mill A, each of the modules contain hydrolyzers, fermenters/acidogens, acetogens, and methanogens. The modular organization is somewhat different than at the other two mills. Modules B1 and B2 are associated with each other because there are six ASVs that are present in both modules. Also, module B1, while containing 57 ASVs, was by far the largest module that was found in any of the reactors. Module B1 is positively correlated to the COD removal efficiency, and negatively correlated to the VFA / alkalinity ratio, and the hydrogen sulfide load in the biogas. On the other hand, module B3 is positively correlated to the hydrogen sulfide load, and negatively correlated to the COD removal efficiency (Figure 5, Table S17). Similar to mill A, one of the modules at mill B (B3) is predominant under conditions characterized by environmental stress and deteriorated treatment performance. In contrast, module B1 thrives when the treatment performance is high, and conditions are more stable. The latter conditions involve low loads of the anaerobic inhibitor sulfide, a low VFA/alkalinity ratio, and a high COD removal efficiency. Numerous ASVs of *Bathyarchaeia* and the fermenters *Dysgonomonadaceae* and *Sphaerochaeta* are part of module B1 and associated with high treatment performance.

**Figure 5.**
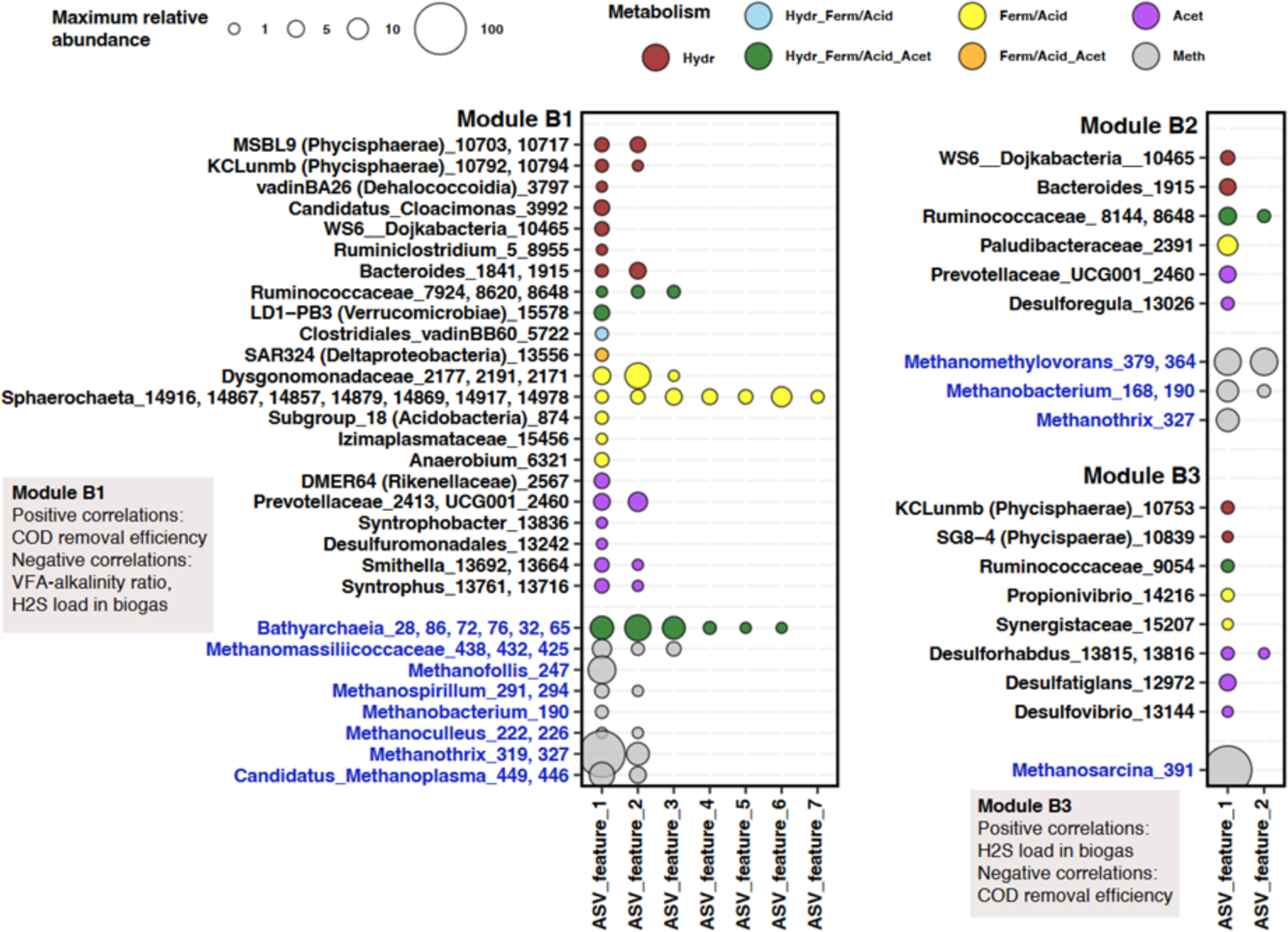
Modules in digester 1 of mill B and correlations with operating parameters. For each module ASVs with the same taxonomic classification are shown on the same line. Black names are bacteria; blue are archaea. Numbers after taxonomy label refers to the ASV feature id in Tables S5-10. Abundances of up to 7 ASVs with the same taxonomy is shown as circles where size refers to relative abundance of the ASV (bacteria and archaea normalized separately). In Module B1 *LD1-PB3* (Verrucomicrobiae) and *WS6_Dojkabacteria* are negatively correlated with COD removal efficiency, and *LD1-PB3* is positively correlated with the VFA-alkalinity ratio, which differs from all other ASVs in the module (Table S17). Hydr – Hydrolysis, Ferm – Fermentation, Acid – Acidogenesis, Acet – Acetogenesis, Meth – Methanogenesis.

Module B3 contains several sulfur and sulfate reducers (ASVs of *Desulforhabdus, Desulfovibrio, Desulfatiglans*) that increase in abundance with increasing sulfide content in the biogas. Several hydrolyzers and fermenters in module B3 cooperate with these sulfur/sulfate reducers which are capable of outcompeting other acetogens and most methanogens in terms of utilizing the produced VFAs as electron acceptors.^51^

Interestingly, module B3 shares ASVs with module A2, which also increased with increasing sulfide content, including *Propionivibrio_*14228*, Paludibacteraceae_*2388*, Lentimicrobiaceae_2836* and *Methanothrix_349* (Figures 4, 5). There are no significant correlations between module 2 and operating parameters. Although the vast majority of high abundance organisms in mill B are organised within these three modules, a few are not. These are *Methanomethylophilaceae*_455 (94%), *Methanomethylovorans_*376 (13%), *Bacteroides*_1935 (15%), *Paludibacter*_2353 (14%), *Sphaerochaeta*_14956 (13%) (and 14944 (6%), 14990 (5%)), and *Prevotella*_7_2437 (7%). Similar to mill A, these organisms are likely less dependent on other organisms and/or changing environmental conditions.

#### 3.2.3 Modules in mill C

The microbial community in the digester at mill C was found to be organized into three modules (Figure 6). Each of the organisms are uniquely assigned to only one module, except for *Clostridiales*_vadinBB60_group_5714 which is included in module C2 and C3. Like the other mills, each of the modules at mill C contains organisms that cover all stages of anaerobic digestion. Modules in the digester at mill C are correlated to similar operating parameters as in the other two mills. Module C1 is positively correlated to the COD removal efficiency and the digester pH, and negatively correlated to the VFA/alkalinity ratio. Module C3 is positively correlated to the VFA/alkalinity ratio, and negatively correlated to the COD removal efficiency and the digester pH (Table S18). Correlations between biogas sulfide concentrations and ASVs in modules C1 and C3 lack significance, likely due to the small number of correlation pairs resulting from substantial missing data (Table S15). Module C3 is more abundant when the community is affected by environmental stress, whereas module C1 prefers operation at stable conditions. Module C3 therefore likely serves a similar purpose as modules A2 and B3 as they all increase in abundance in response to process inhibition, indicated by an elevated sulfide concentrations and/or a high VFA-alkalinity ratio. Modules A1, B1, and C1 increase when sulfide levels or the VFA-alkalinity ratio are low, and the anaerobic treatment performance is high. These results highlight the detrimental impact that high sulfide concentrations have on the anaerobic microbial community and the reactor operating conditions.

**Figure 6.**
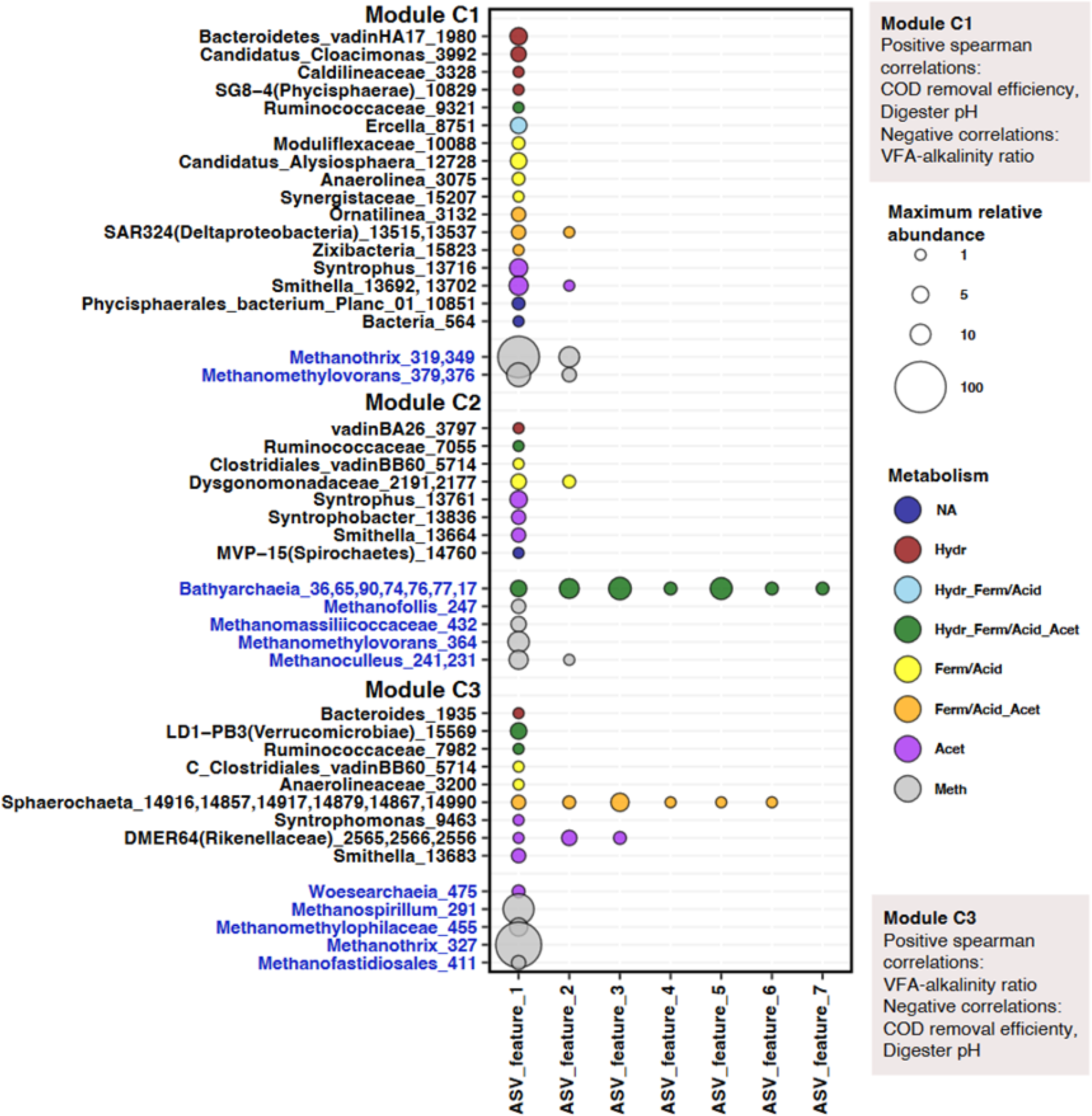
Modules in the digester at mill C and correlations with operating parameters. For each module ASVs with the same taxonomic classification are shown on the same line. Black names are bacteria; blue are archaea. Numbers after taxonomy label refers to the ASV feature id in Table S5-10. Abundances of up to 7 ASVs with the same taxonomy is shown as circles where size refers to relative abundance of the ASV (bacteria and archaea normalized separately). Hydr – Hydrolysis, Ferm – Fermentation, Acid – Acidogenesis, Acet – Acetogenesis, Meth – Methanogenesis.

Module C2 contains several ASVs of *Bathyarchaeia* and *Dysgonomonadaceae* that are positively correlated to the concentration of color in the final effluent (Table S18). According to the study by Yousefian,^52^ color in pulp mill wastewater consists of 50 – 90% of lignin derived compounds. As previously mentioned *Bathyarchaeia* has been associated with lignin degradation. Therefore, a higher color content in the wastewater may induce the growth of these archaea. In numerous previous studies, members of the family *Dysgonomonadaceae* have been found in the guts of insects that feed on lignocellulosic rich organic matter and in anaerobic digesters with lignocellulosic material as the main substrate.^53^, and refs therein As in the two other mills, several high-abundance ASVs could not be assigned to any of the three modules. These ASVs are *Paludibacter*_2351 (19%), *Prolixibacteraceae_*2527 (18%), and Subgroup_18 (Acidobacteria)_874 (5%).

### 3.3 Three-way associations – mediation by a third factor

Liquid association (LA) analysis as part of the LSA software enables the identification of three-feature associations.^16^ This analysis examines how co-occurrence might be mediated by a third biological or environmental variable. Liquid associations between three ASVs, as shown in Figure 7, are relationships where a non-existent or weak correlation between two ASVs is strengthened by the abundance of a third ASV. Figure 7 only includes high abundance (>5%) ASVs in the cases of mills B and C. Compared to the other mills, mill A displayed fewer instances of such three-way relationships. As a result, some triples were included in the figure that include ASVs with lower maximum abundance (<5%) (Figure 7 – upper left). In the mill digesters, the vast majority of the three-way associations consist of acidogenic fermenters and/or archaeal organisms. Archaea involved in these relationships are *Bathyarchaeia, Methanothrix, Methanomethylovorans, Methanomassiliicoccaceae, Methanomethylophilaceae, Methanobacterium* and *Candidatus_Methanoplasma*, and the acidogens/fermenters belong to *Anaerolineaceae*, *Lentimicrobiaceae*, *Sphaerochaeta*, *Desulfobulbus, Dysgonomonadaceae, Anaerolineaceae*, *Proteiniphilum*, and *Subgroup18 (*phylum Acidobacteria). Acidogenesis has been referred to as the product-determining step in anaerobic digestion because it involves numerous symbiotic relationships within the microbial community and disruptions in the acidogenesis step has the potential to cause process inhibition.^54^ Some of the identified three-way associations may refer to symbiotic relationships involving fermenters/acidogens and archaea. All of the triples involve either two mediated organisms within the same module, or at least one of the mediated organisms not belonging to any module. Some of the mediating ASVs are part of a different module which may refer to competitive relationships between mediating and mediated ASVs.

**Figure 7.**
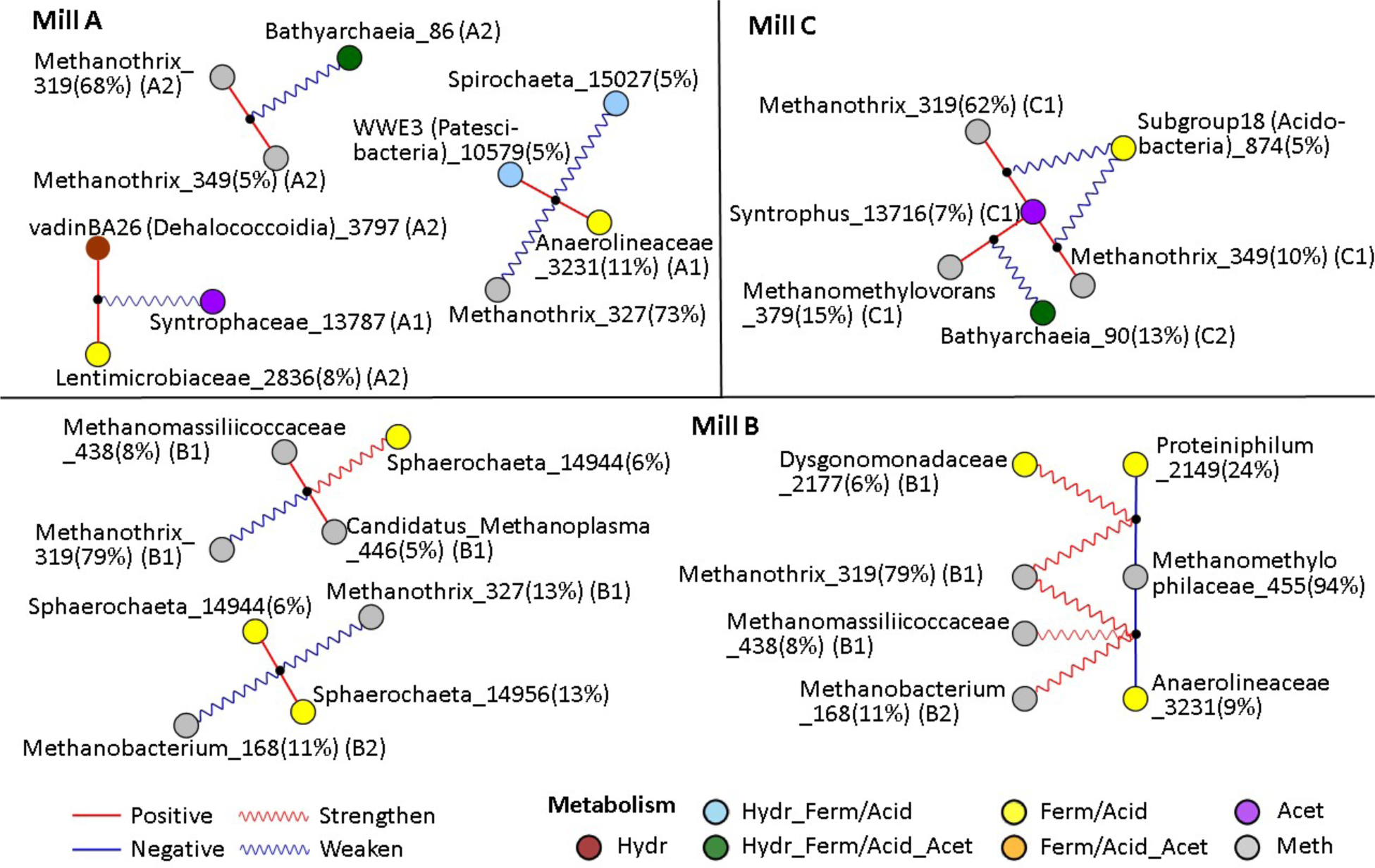
Three-feature associations between ASVs. Wavy lines connect enabling third party to pairs of ASVs connected by straight lines. Red/blue straight lines: positive/negative correlations. Red wavy lines: an increasing abundance of the mediating ASV increases the strength of the correlation (positive or negative) between two ASVs. Conversely, a blue wavy line means that the 3^rd^ party decreases the strength of the correlation between the other two ASVs. For example, in mill A, increasing *Bathyarchaeia_*86 abundance decreases the strength of the positive correlation between *Methanothrix*_319 and *Methanothrix*_349. Module designation is the same as in section 3.2. Hydr – Hydrolysis, Ferm – Fermentation, Acid – Acidogenesis, Acet – Acetogenesis, Meth – Methanogenesis.

In numerous cases, an increase in the abundance of the mediating organism weakens the positive correlation between the two other ASVs (Figure 7, blue wavy lines, red straight lines). These cases involve a declining association of two organisms that previously cooperated or were adapted to the same environment, by a third mediating organism. In mill C, the growing presence of *Subgroup18* from the phylum Acidobacteria might disrupt the symbiotic relationship between *Methanothrix_*319 and *Syntrophus_*13716. This disruption could occur if Subgroup18 forms a preferred association with one of the two ASVs (Figure 7). Conversely, at mill B, there are a few instances where an increase in the abundance of the mediating ASV amplifies the negative correlation between two ASVs (Figure 7, red wavy lines, blue straight lines). This may indicate an increasing competition between the two mediated ASVs, possibly leading to mutual exclusion, which is mediated by the presence of a third ASV.

It is worth noting that there is a scarcity of scenarios where an increase in the abundance of a mediator enhances a positive correlation between two other ASVs. This finding seems to support Palmer and Foster’s ^55^ study, emphasizing the rarity of cooperation in microbial relationships.

Liquid associations were also identified including operating parameters as mediating features (Figure S6). These associations may be interpreted as correlations between ASVs that increase or decrease in strength as a response to a change in operating conditions. As an example from the digester at mill B, an increasing VFA-alkalinity ratio, indicating process deterioration, is related to a stronger correlation between the two fermenters *Synergistaceae* and *Ruminococcaceae*.^56,57^ Also, a high daily load of the anaerobic inhibitor hydrogen sulfide in the biogas increases the correlation between two ASVs of *Proteiniphilum* and *KCLunmb (Phycisphaerae)*. Both organisms have been previously linked to fermentation and hydrolysis,^41,58^ respectively. Elevated concentrations of inhibitory compounds and the associated process deterioration seem to cause an increased cooperation or dependence between fermenting, acidogenic and hydrolytic organisms to counteract the effects of the environmental stress. At mill A, a rise in the level of sulfite in the reactor influent amplifies two negative correlations and weakens one positive correlation among the three organisms *Candidatus_Endomicrobium_*4288*, Anaerolinea_*3076, and *Georgenia_*1277. The associations suggest the existence of competitive dynamics between mediated ASVs as well as between mediated and mediating ASVs. Beyond these three-way associations there probably also exist numerous relationships between four and more organisms, however these relationships are likely impossible to identify with the currently available tools.^13^ These relationships can help inform and eventually improve the annotation of these poorly understood taxa.

### 3.4 Lagged parameter relationships

Because there is a delay between changes in wastewater treatment operation and the responding change in microbial abundance, lagged operating parameters were generated and correlations with microorganism abundance were calculated using LSA. To minimize false discovery rates and identify meaningful time lags between autocorrelated time series we applied a significance level at p<0.001 and q<0.05. The oxidation-reduction potential in the reactor feed basin, and the associated sulfide concentration in the reactor influent, showed highest lagged correlations with the abundance of numerous microorganisms in mill A. As shown in section 3.2, this parameter is correlated to ASVs from the modules A1 and A2. Figure 8 shows correlations between sulfide concentrations, lagged by 0 to 5 days, and 11 ASVs. Higher sulfide concentrations (lower ORP) negatively affect the abundance of ASVs from *Anaerolineaceae* and *Methanobacterium*, several days later. The decline of *Methanobacterium* abundance may be due to reduced conditions (low ORP) that promote the growth of strictly anaerobic homoacetogenic bacteria that outcompete hydrogenotrophic methanogens for hydrogen.^59^ Higher sulfide concentrations are also correlated to an increasing abundance of ASVs from for instance *Bacteroidetes_vadinHA17, Lentimicrobiaceae, Methanothrix, Paludibacter,* and *Propionivibrio*, several days later. These organisms may be more resistant to environmental stress caused by high sulfide levels. Generally, the strongest correlations were found between ASV abundance and sulfide concentrations that were lagged in time by 2 – 3 days (Figure 8). Therefore, this period likely corresponds to the average response time between the change in digester operation and the change in organism abundance at mill A.

**Figure 8.**
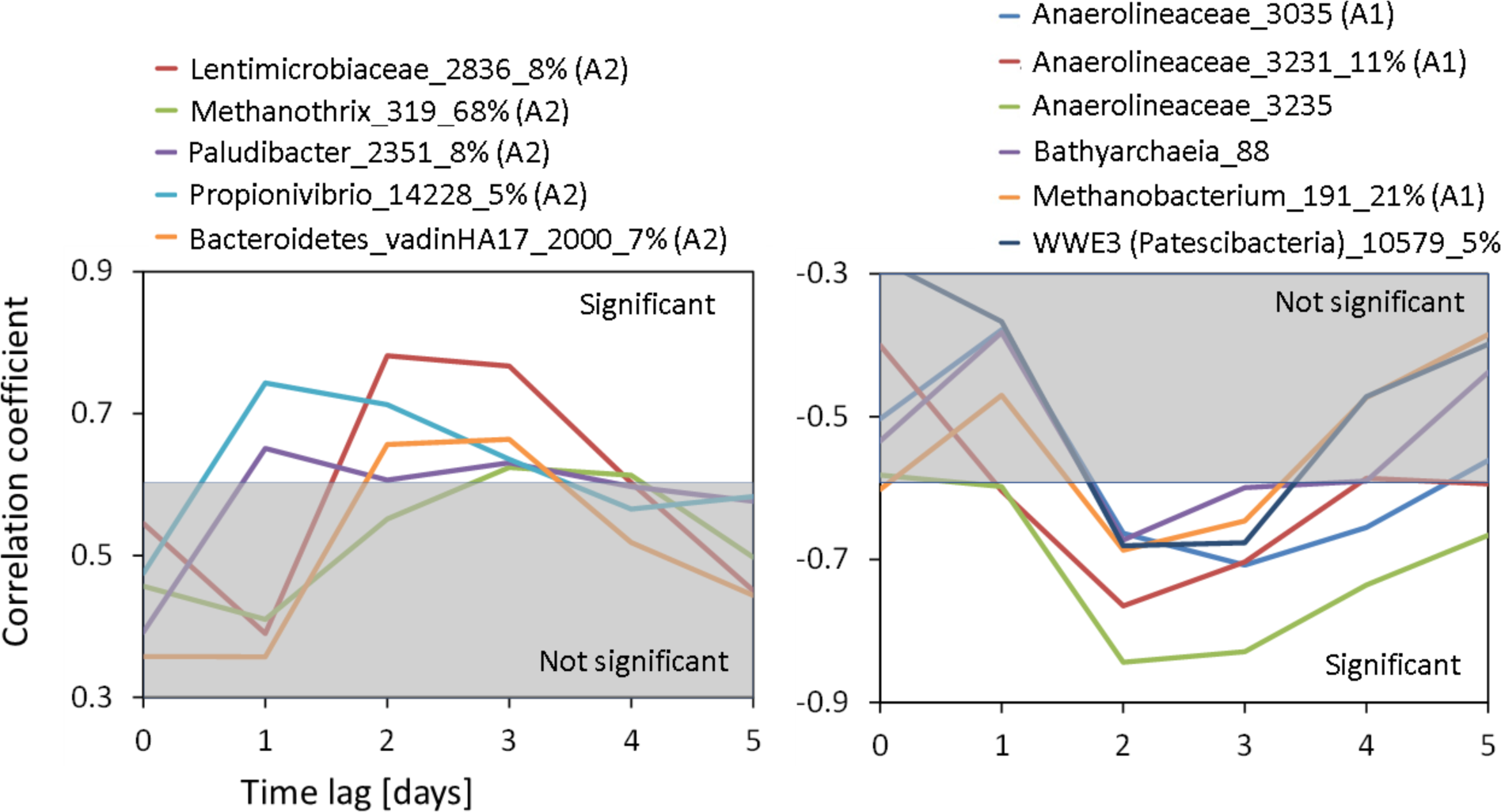
Correlations between lagged sulfide concentrations in the reactor feed basin and microorganism abundance at mill A. Values outside the shaded areas are significant at the 99.9% level. The lag of 0 days means that no time lag was applied. Module designation is the same as in section 3.2.

At mill B there are several correlations between lagged operating parameters and individual microorganisms at p < 0.001. Highest correlations were found between various ASVs from the modules B1 and B3 and lagged versions of the parameter daily hydrogen sulfide (H_2_S) load in the biogas (Figure 9). Other time-lagged operating parameters such as the daily sulfur load in the digester influent and the organic feed load are also significantly correlated to the organism abundance. An increase in sulfur load expectedly increases the abundance of several sulfur and sulfate reducers (Figure S8). Several organisms are negatively affected by environmental stress conditions caused by a high sulfide load and a high COD load. These are ASVs from *Dysgonomonadaceae, Izimaplasmataceae,* and *Smithella* (Figures 9, S9).

**Figure 9.**
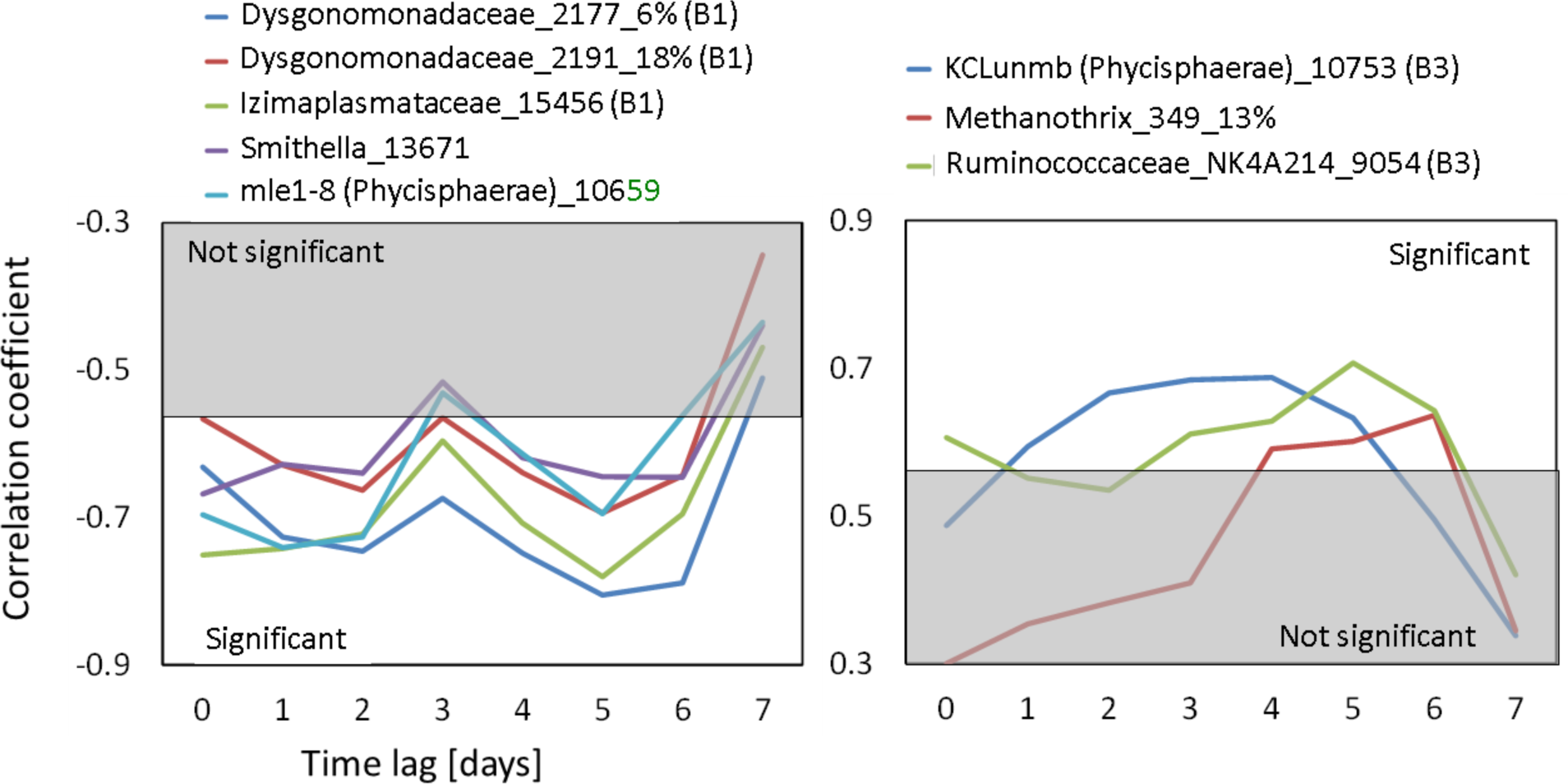
Correlations between the lagged daily hydrogen sulfide (H_2_S) load in the biogas and microorganism abundance at mill B. Values outside the shaded areas are significant at the 99.9% level. Module designation is the same as in section 3.2.

At mill B, the response time between the changing operating conditions and changes in organism abundance ranges between 2 and 6 days and therefore is somewhat longer compared to mills A and C. The reason might be that the washout of ASVs along with the digester effluent at mill B is slower than at the other two mills. This is indicated by a relatively long hydraulic retention time, and relatively small total suspended solids (TSS) concentrations in the effluent, which were interrupted by short-term spikes of very high TSS concentrations. This may have led to an increase in the response time, and a widening of the curves in Figures 9, S8, S9. Also, and as previously mentioned, statistical artefacts such as autocorrelation of the time series can distort and shift the true lag time.

At mill C, besides the treatment performance parameters, the parameter whose lagged versions showing the strongest correlations with the abundance of ASVs is the pH of the digester. This parameter is strongly associated with two of the identified modules, C1 and C3 (Figures 6, 10).

**Figure 10.**
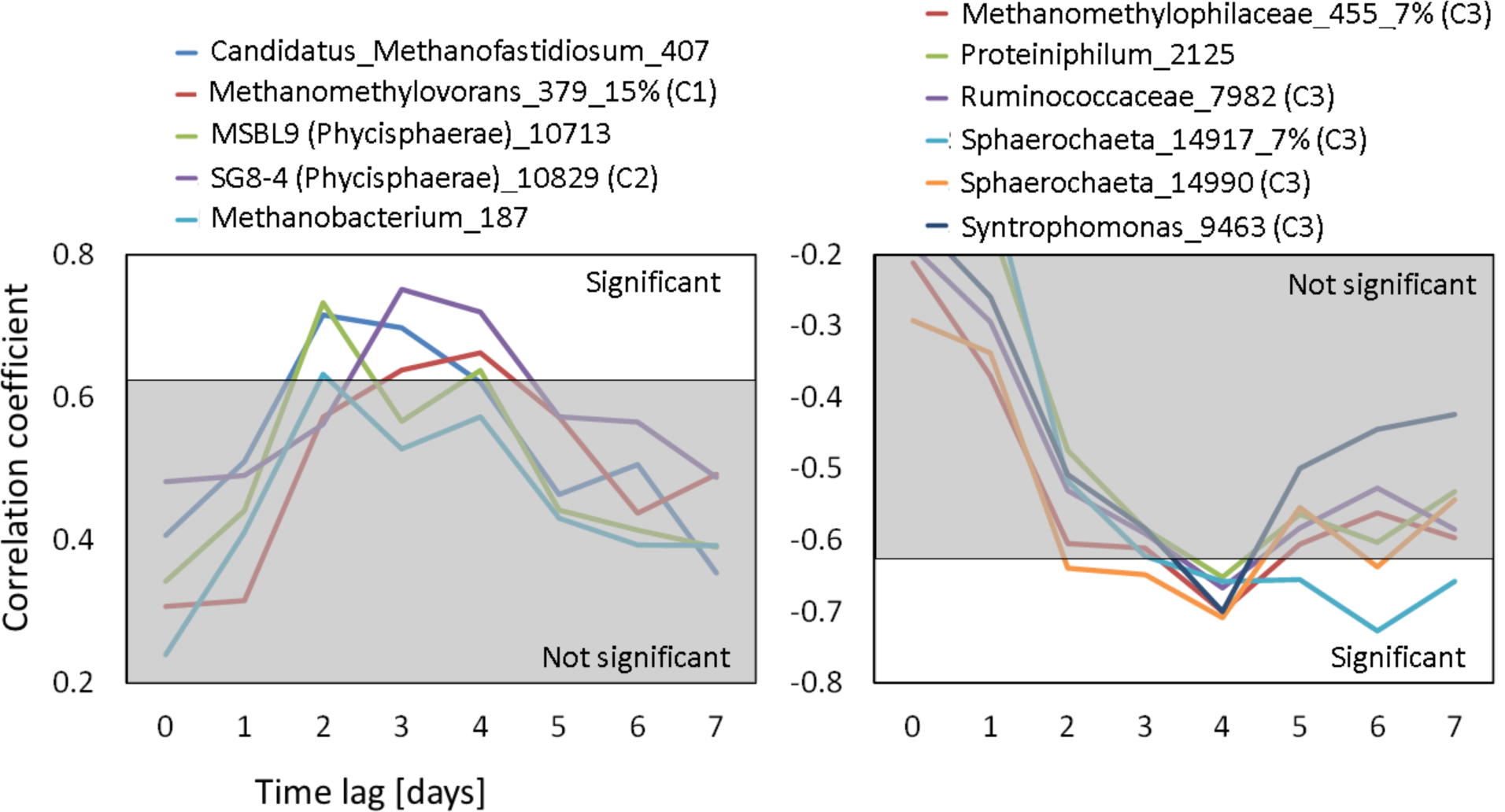
Correlations between the lagged parameter digester pH and various microorganisms at mill C. Values outside the shaded areas are significant at the 99.9% level. Module designation is the same as in section 3.2.

Many correlations between operating parameters and ASV abundance would not have been identified without using lagged parameters, because correlations with operating parameters that are not lagged in time (lag of 0 days) are usually smaller and often not significant.

### 3.5 Digester upset events and the microbial community response

The digesters at mill A experienced only one notable process disturbance during the investigation, which was caused by an annual mill shut down. The upset period, including the shut down, occurred between May 08 and May 24, 2017 (Figure 11). The VFA-alkalinity ratio (based on concentrations in meq/l) was 0.27 when it peaked on May 14. The average ratio during the study period was 0.06. The lowest COD removal efficiency during that time was 44% (average: 61%). The biomass samples collected on Apr 22, May 19, May 31, and Jun 30 were used to investigate the change in the abundance of individual organisms as a result of the upset. Microorganisms whose relative abundance more than doubled include ASVs from the VFA and long-chain fatty acid degrading syntrophic bacteria *Syntrophobacter*, *Syntrophaceae*, and *Synergistaceae*, as well as several ASVs from *Anaerolineaceae* (Figure S10). Accelerated growth of these acid degrading bacteria may have prevented an acidification of the digesters. During the upset the pH was kept at around 6.8 and therefore, did not decrease significantly. Interestingly, the ASV *Methanothrix*_319 increased in abundance and responded to upset conditions in a similar way in all three mills (Figure 11). Also, under stable conditions several *Anaerolineaceae* ASVs are strictly associated with hydrogenotrophic methanogenesis (Figure 4). However, during the upset period these ASVs seem to thrive at similar conditions as the acetoclastic *Methanothrix*_319, suggesting a possible mutualistic relationship in this environment, and pointing to the flexibility of members of the *Anaerolineaceae* family.

**Figure 11.**
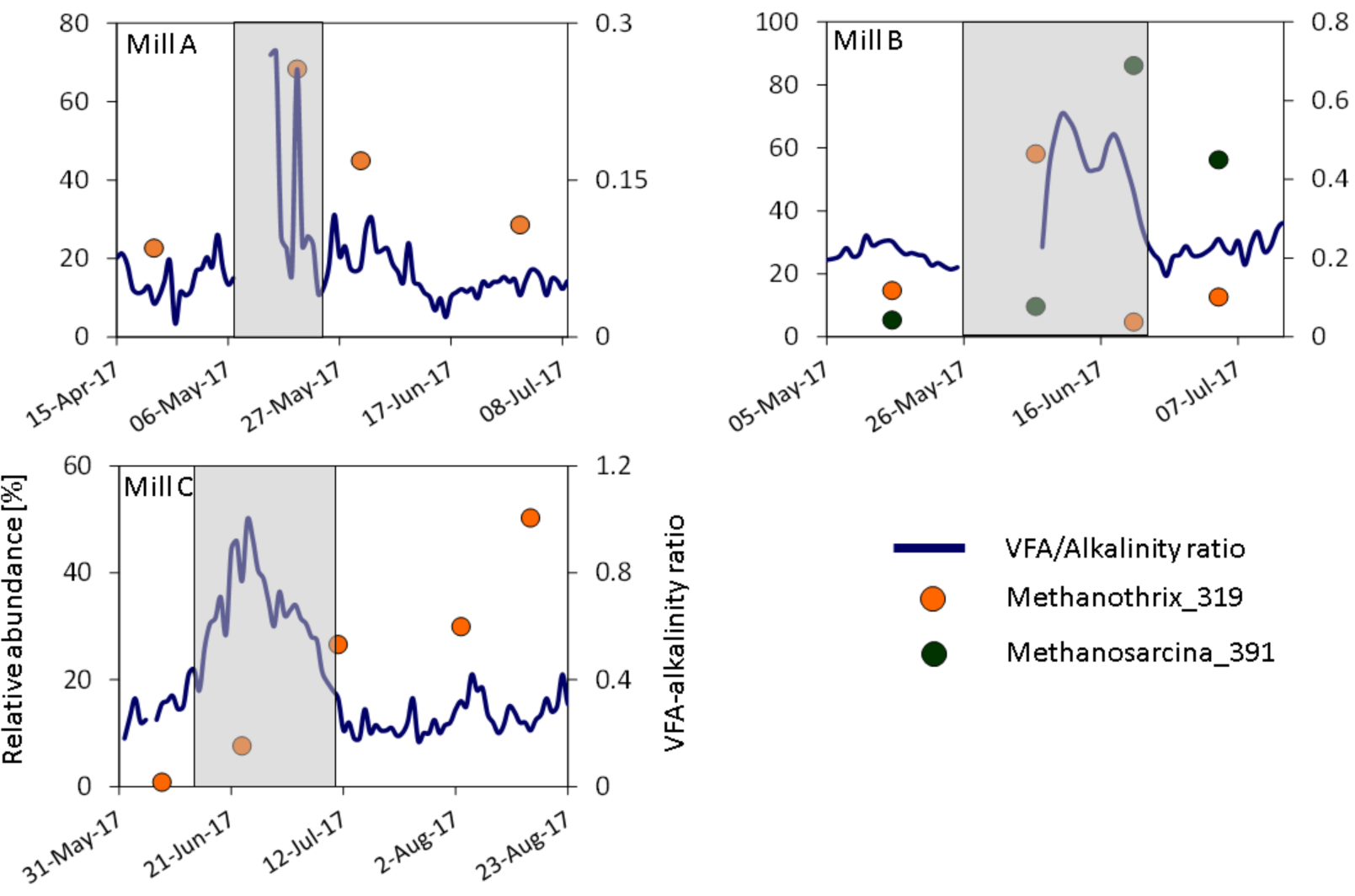
VFA-alkalinity ratio, and abundance of *Methanothrix_*319 and *Methanosarcina_*391 during upset periods at the three mills. The gap in the line graphs is due to the annual mill shutdown periods in mills A and B. Shaded area: upset periods. Please note that mill A used a different titration method to determine the VFA-alkalinity ratio than the other two mills.

At mill B, the most significant period of process impairment occurred due to an annual mill shut down during which the digesters were out of operation for ten days. The upset period started with the shut down and lasted from May 26 to June 22, 2017. The VFA-alkalinity ratio was 0.57 (based on mg/l) when it peaked on June 10 (average ratio: 0.26). At that time the COD removal efficiency was 37% (average: 61%). The lowest pH during this period was 6.9 and therefore did also not decrease notably. The ASV abundance in the samples collected on May 15, June 06, June 21, and July 04 were used to investigate the impact of the process impairment. The first and the last of these four samples were collected during normal operating conditions. Like mill A, the community in mill B experienced an increase in acetoclastic methanogens which partially replaced hydrogenotrophic and methylotrophic methanogens. Within about two weeks (June 06 - 21) a *Methanosarcina* ASV replaced *Methanothrix_*319 and became the dominant archaea (Figure 11). *Methanosarcina spp*. have been shown to be resistant to various types of anaerobic stressors such as high ammonium, sodium and acetate concentrations as well as sudden pH changes.^60^ Other high-abundance organisms that increased in relative abundance in response to the upset conditions were ASVs from *Proteiniphilum*, *Desulfatiglans, Candidatus Cloacimonas, KCLunmb (Phycisphaerae)*, *Ruminococcaceae*, and *Anaerolineaceae* (Figure S11). These bacteria have been associated with hydrolysis, fermentation, acidogenesis and sulfate reduction. Similar to mill A, members of the *Anaerolineaceae* family increased in relative abundance, once again corroborating the key role that these organisms play during anaerobic wastewater treatment.

At mill C, the only major period of process deterioration lasted from June 15 to July 10, 2017. The VFA-alkalinity ratio peaked on June 24 when it was 1.0 (based on mg/l) (average: 0.31), and the COD removal efficiency at this time was 15% (average: 67%). Like in the other two mills a notable pH decrease was averted. This was accomplished by temporarily decreasing the organic loading rate and diverting parts of the wastewater directly into aerobic treatment, while bypassing the digester. At mill C the reason for the upset was a combination of organic overload and nitrogen deficiency. The samples collected on June 08, June 23, July 11, August 03, and August 16 were used to assess the effects of the process impairment. The sample from June 23 almost coincides with the peak of the upset period. Therefore, the community in this sample should strongly reflect the impact of the process upset. As in the other mills *Methanothrix_*319 increased in relative abundance during the upset. However, the dominance of this ASV was not obvious until July 11 when the following sample was collected. Other ASVs that increased in abundance include the hydrolyzers and fermenters *Prolixibacteraceae, Bathyarchaeia*, and *Sphaerochaeta* (Figure S12).

Considering the differences between the digesters in the three mills in terms of reactor type, wastewater composition, original inoculum, and the number of years of operation, it is remarkable that the same ASV, *Methanothrix_*319, grew and became the dominant archaeal organism as a response to process upset events. The maximum relative abundance (based on the archaeal population) of *Methanothrix_*319 was 68% at mill A, 58% at mill B, and 27% at mill C. The latter percentage would have been notably higher if there were not several high-abundance *Bathyarchaeia* present in the community at this time. The physiology of Methanothrix_319 needs to be explored further to better understand why the organisms represented by this ASV became so enriched in these periods of upset. This ASV may represent closely related organisms with highly beneficial traits that may even be candidates for bioaugmentation or could represent a group of highly resilient organisms that survives upset periods while other microbes do not. Shotgun metagenome sequencing has been completed on six selected samples from these three mills for further investigating this and other questions and the data are available under NCBI project PRJNA916529.

## 4 Conclusions

Amplicon sequencing and operating data collected over 1.5 years from anaerobic wastewater treatment reactors at three pulp and paper mills were used to investigate the structure of the microbial community and its relationships to operating parameters.

Two to three relatively stable biologically functional groups or modules emerged from the analysis of the microbial communities in the digesters. Each contained hydrolyzers, fermenters/acidogens, acetogens, and methanogens. The modules were correlated to various operating parameters such as the concentration of sulfide and the pH in the digester, as well as anaerobic treatment performance parameters. Also, the modules show an antagonistic response to some of these parameters. For instance, an improvement in anaerobic wastewater treatment performance is associated with an increasing abundance of one module, and a decreasing abundance of a second module. Therefore, these functional modules balance the anaerobic digestion process in response to the varying operating conditions.

Methane generation in all digesters was dominated by acetoclastic methanogens, which represent more than half of the archaeal population. The study also enhances insight into *Bathyarchaeia*, which were present in all three mills at relatively high abundance, potentially linked to the elevated lignin content in the mill wastewater. In all digesters *Bathyarchaeia* accounted for 10% to 20% of the total archaeal community. Also, most ASVs from *Bathyarchaeia* were positively correlated to hydrogenotrophic and/or methylotrophic methanogenesis.

The investigation of the correlations using time lagged parameters revealed that the response time between the change of an operating parameter and the change in relative organism abundance is usually between two to four days. This response time may be related to the growth rates of anaerobic microorganisms, the hydraulic retention time, and the biomass washout rates in the individual digesters.

Upset conditions, as a result of plant shut down events or organic overload, caused a drastic change of the microbial community composition. As a response to these conditions, the ASV *Methanothrix_*319 increased in abundance and dominated the archaeal population in all three mills. The closely related organisms associated with this ASV may be considered for growing bioaugmentation cultures to be applied in cases of process upsets and should be further investigated.

The findings of this study are expected to have relevance for anaerobic wastewater treatment across various industries and municipalities, because of the similarities observed in the microbial composition, trophic stages of anaerobic digestion, and operating conditions among different digesters.

## Supporting information

Supplemental large tables

## Associated Content

Additional data including correlations between modules and operating parameters, lagged parameter correlations, relative abundances of ASVs, normalized daily operating parameters, and modular assignment values are available (Supporting Information and Supporting Tables). All sequencing data related to this project are deposited at NCBI under project PRJNA916529.

## Author Contributions

TM implemented the correlation and module analyses and wrote the paper. MIY conducted the DNA extraction and the analysis of the amplicon sequences, and proofread the manuscript. EAE and CN contributed to the interpretation of the results and proofread the manuscript. EM proofread the manuscript. All authors approved the final manuscript.

## Notes

The authors declare no competing financial interest.

## Acknowledgements

We are indebted to the personnel of the three mills for their help with sampling and consultations. Special thanks go to Shen Guo for helping with the DNA extraction.

