## Supporting Information for "Microbial community organization during anaerobic pulp and paper mill wastewater treatment"

### Contents


| **Material** | | **Page** |
|---|---|---|
| Figure S1 | Heatmap mill B | S2 |
| Figure S2 | Heatmap mill C | S3 |
| Figure S3 | Community network – mill A | S4 |
| Figure S4 | Community network – mill B | S4 |
| Figure S5 | Community network – mill C | S5 |
| Figure S6 | Three-feature associations with operating parameters as mediating factors | S5 |
| Figure S7 | Generation of lagged parameters | S6 |
| Figure S8 | Correlations between lagged sulfur load and microorganisms at mill B | S7 |
| Figure S9 | Correlations between lagged COD load and microorganisms at mill B | S7 |
| Figure S10 | Organisms with an abundance change >100% during the upset period at mill A | S8 |
| Figure S11 | Organisms with an abundance change >100% during the upset period at mill B | S8 |
| Figure S12 | Organisms with an abundance change >100% during the upset period at mill C | S9 |
| Table S1 | Correlations between Bathyarchaeia and hydrogenotrophic and methylotrophic methanogenesis | S10 |
| Tables S2-S4 | In accompanying Excel file – Modular assignment values for each mill | Excel |
| Tables S5-S10 | In accompanying Excel file – ASV taxonomy and relative abundance data for all digesters | Excel |
| Table S11 | In accompanying Excel file - Representative ASV sequences | Excel |
| Table S12 | Amplicon sequencing reads in each QIIME step | Excel |
| Tables S13-S15 | In accompanying Excel file - Daily operating parameters for each mill, normalized to the range [0, 1]. Legend shown on p.S11 for reference | S11& Excel |
| Tables S16-S18 | In accompanying Excel file – Correlations between modules and operating parameters for each mill | Excel |

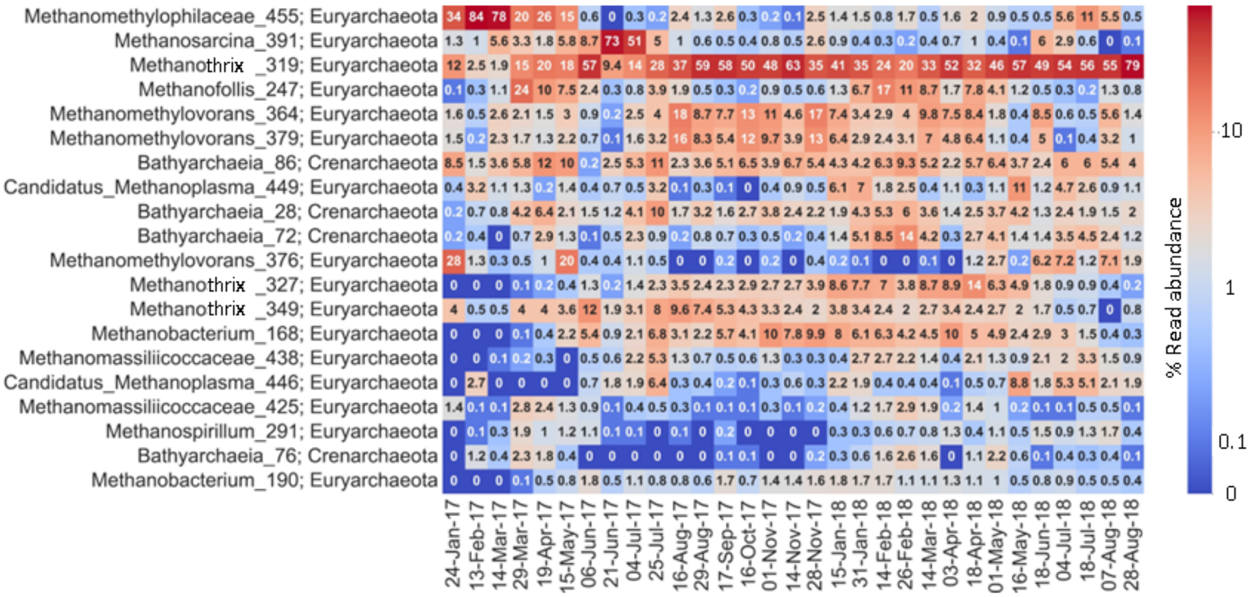


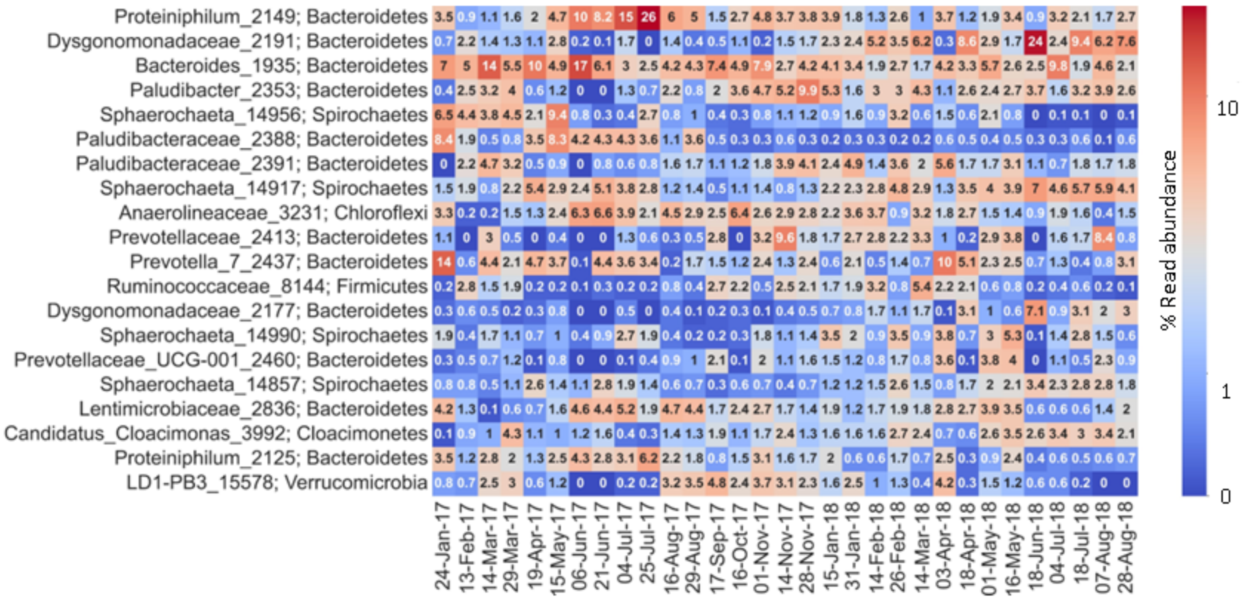


Figure S1 Heatmap with relative abundances of archaea and bacteria, averaged over the three digesters at mill B.

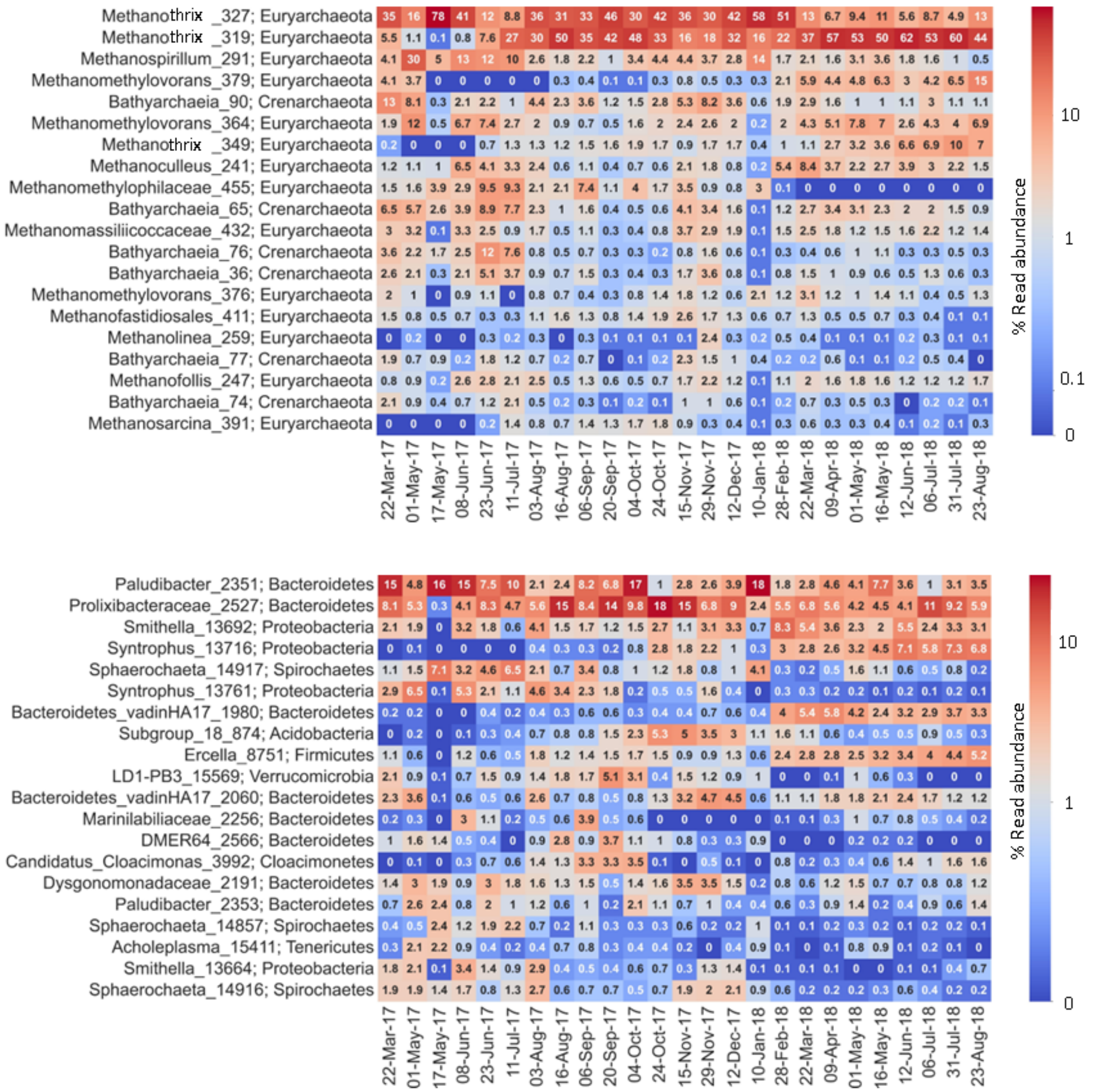


Figure S2 Heatmap with relative abundances of archaea and bacteria in the digester at mill C.

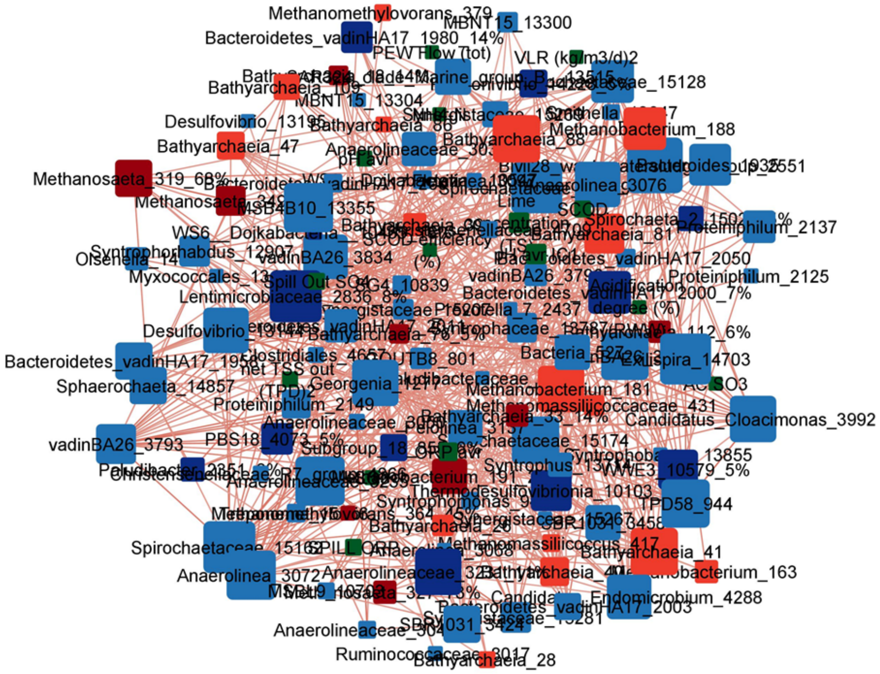


Figure S3 Correlation network of the microbial community in digester 1 at mill A. Dark red: high abundance archaea, bright red: low abundance archaea, dark blue: high abundance bacteria, bright blue: low abundance bacteria, green: operating parameters, larger squares mean a higher node degree, i.e. more significant co-occurrences with other organisms.

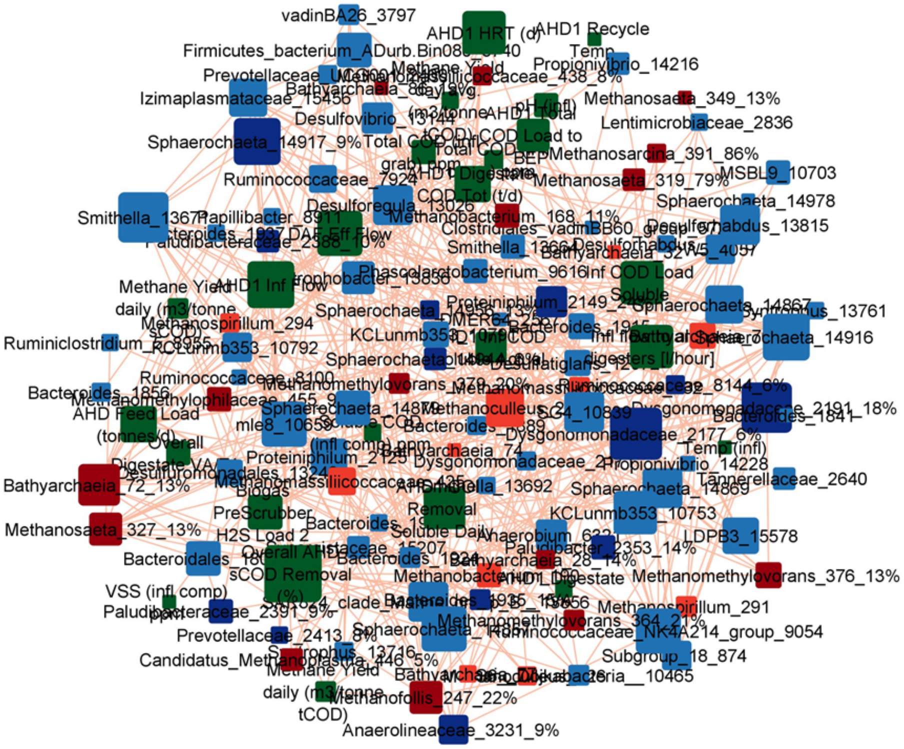


Figure S4 Correlation network of the microbial community in digester 1 at mill B.

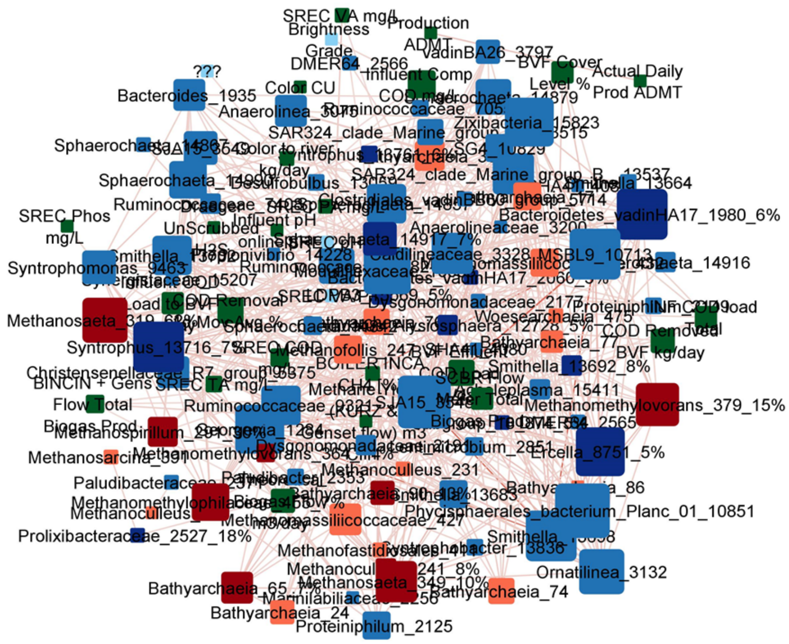


Figure S5 Correlation network of the microbial community at the digester from mill C.

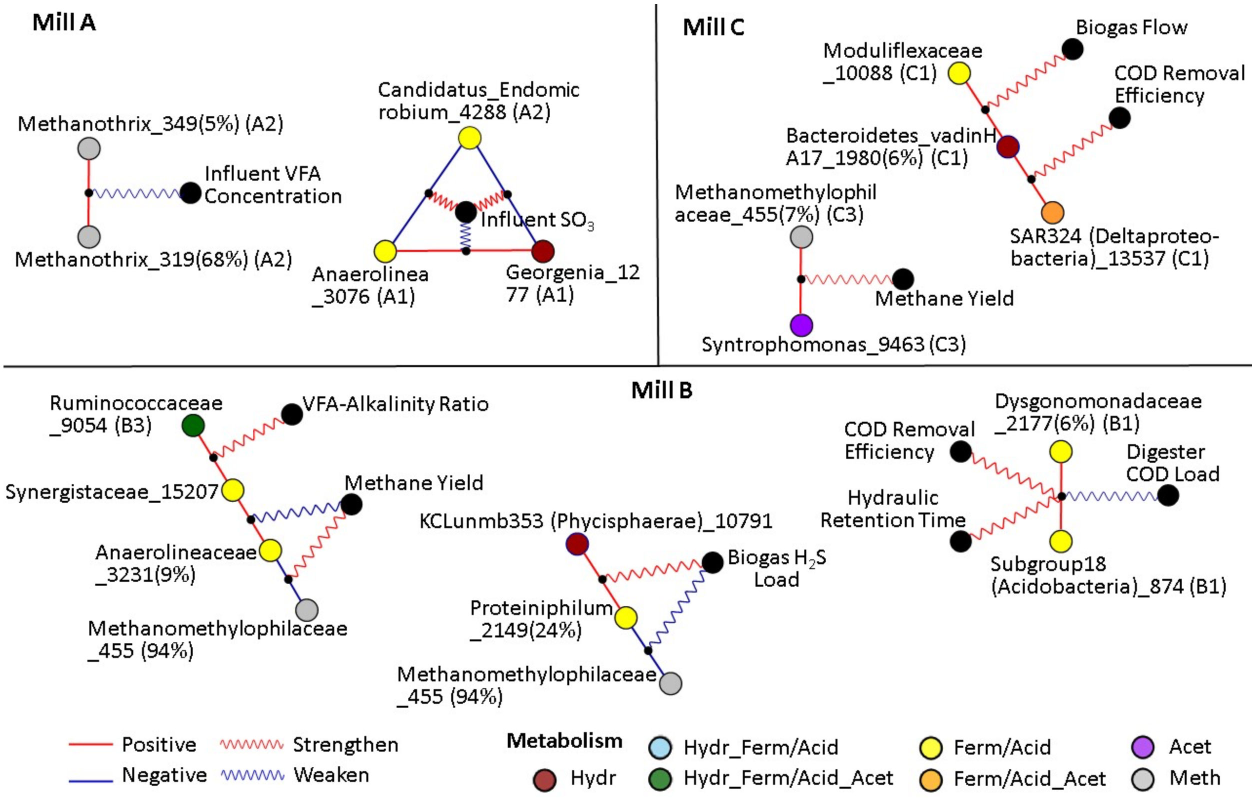


Figure S6 Three-feature associations with operating parameters as mediating features. Red lines: positive correlations, blue lines: negative correlations. Red wavy line means that increasing abundance of mediating variable increases the correlation strength between two ASVs. Blue wavy line is converse. VFA: volatile fatty acids, COD: chemical oxygen demand. Module designation is the same as in section 3.2. Hydr – Hydrolysis, Ferm – Fermentation, Acid – Acidogenesis, Acet – Acetogenesis, Meth – Methanogenesis.

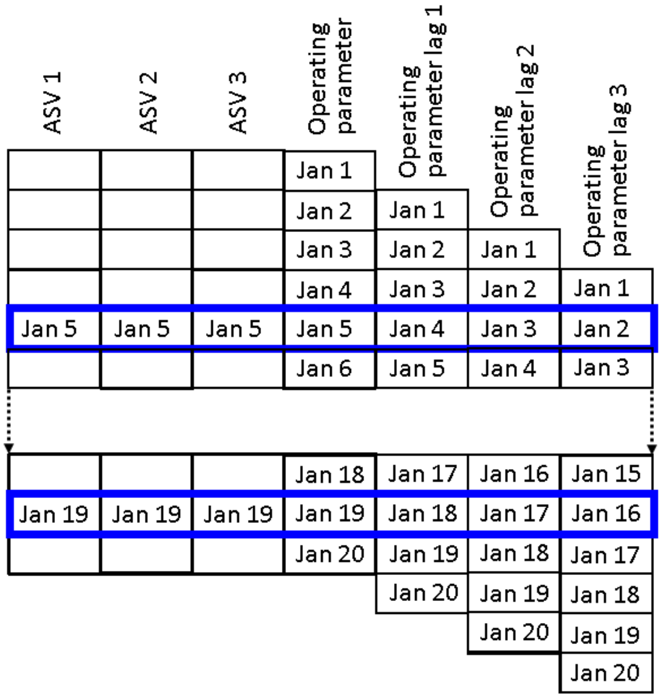


Figure S7 Illustration of methodology to generate lagged parameters for a hypothetical time period (Jan 1–20). This example includes three ASVs each from two samples (Jan 5, 19), and one operating parameter, measured once a day. The operating parameter was shifted forward to create lagged versions of the parameter (lag 1-3). This enables correlating the ASV abundance with the operating parameter values measured one, two, and three days earlier.

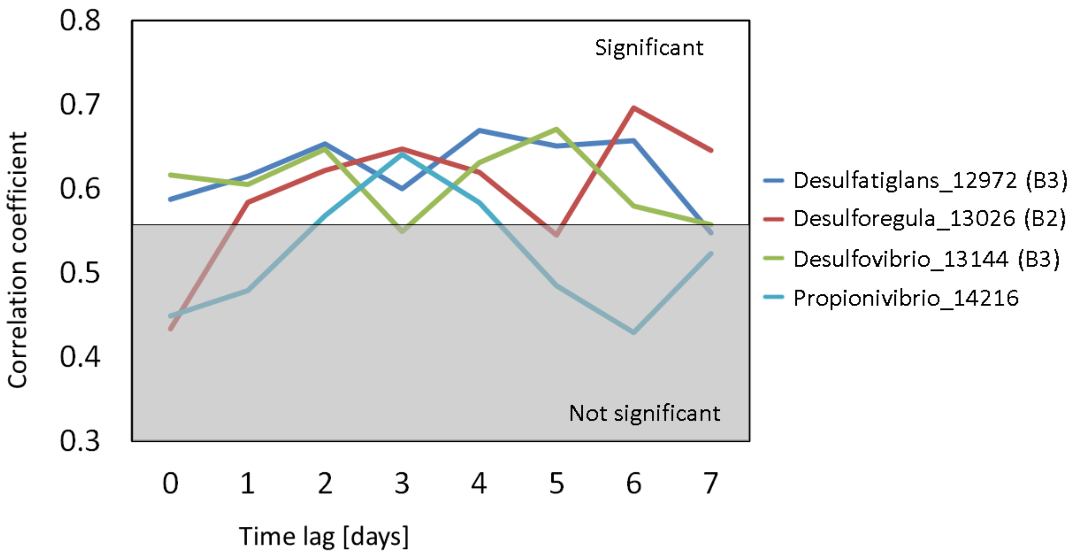


Figure S8 Correlations between lagged sulfur load to the digesters and various microorganisms at mill B.

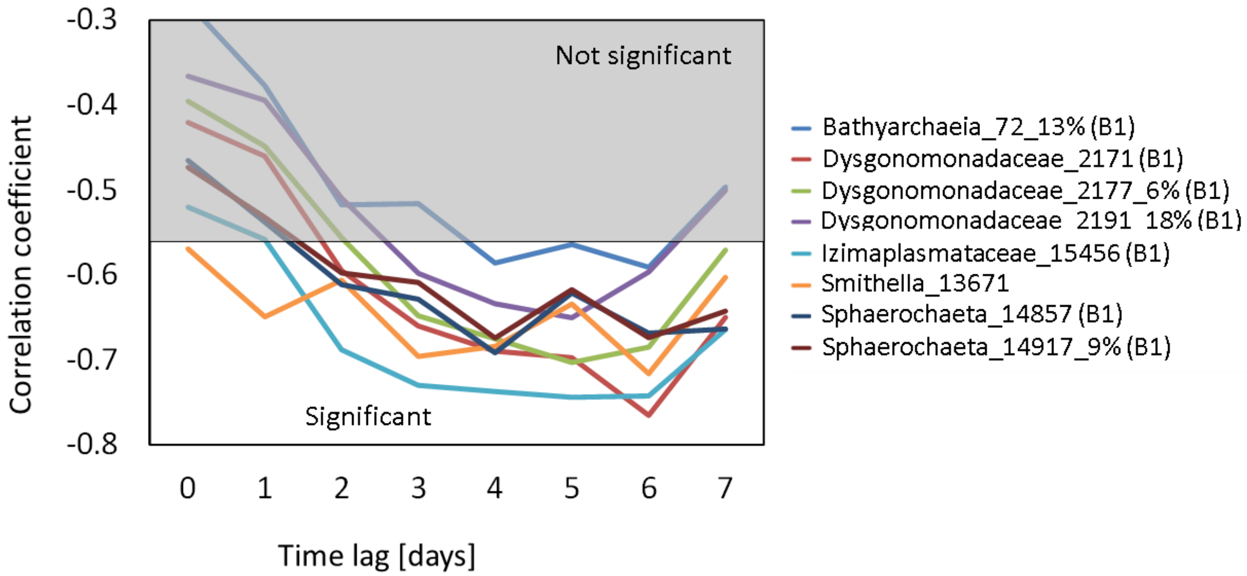


Figure S9 Correlations between lagged digester COD feed load and various microorganisms at mill B.

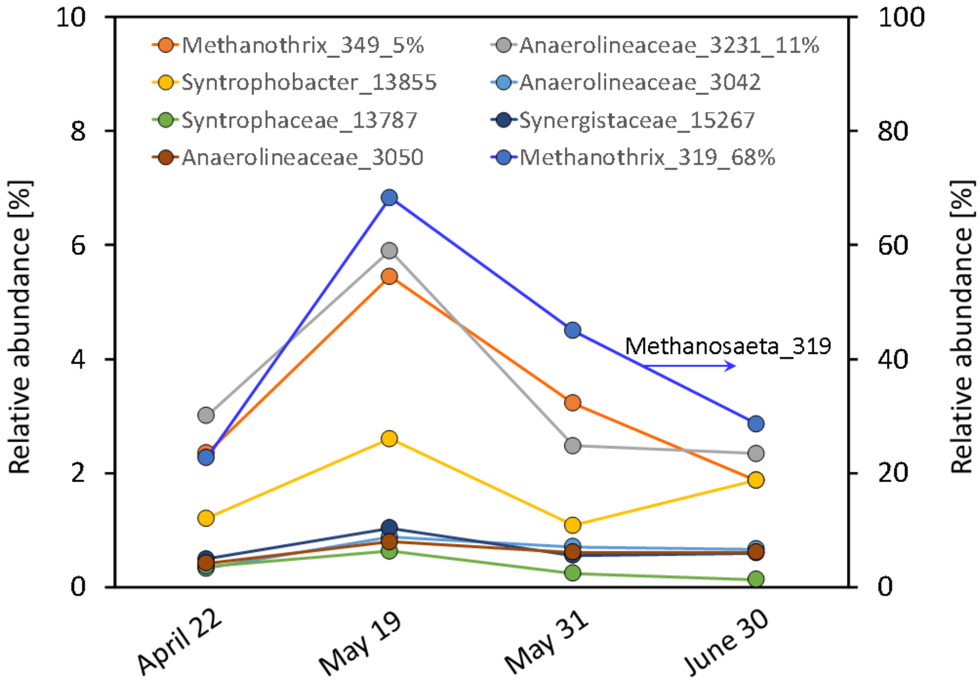


Figure S10 Organisms with an abundance change by >100% during the upset period (May 08-24, 2017) caused by an annual shutdown at mill A.

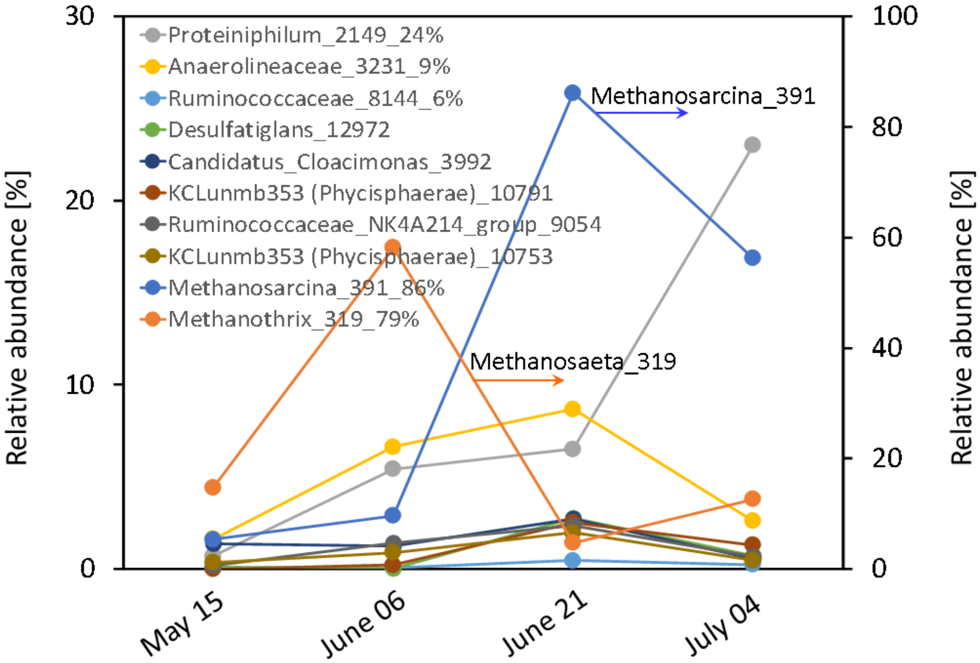


Figure S11 Organisms with an abundance change by >100% during the upset period (May 26 - June 22, 2017) caused by an annual shutdown at mill B.

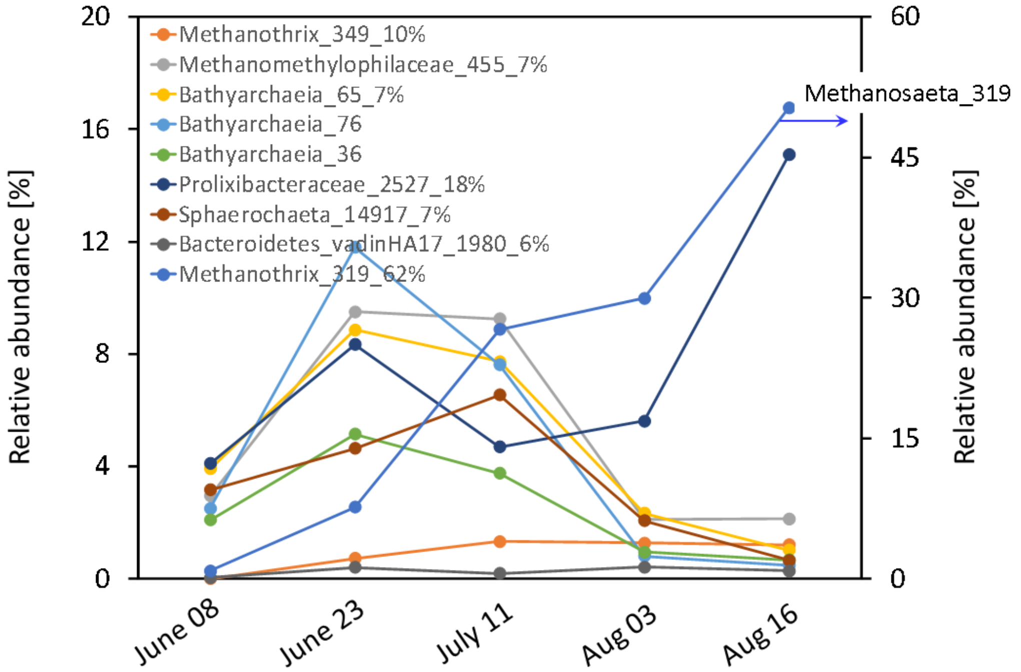


Figure S12 Organisms with an abundance change by >100% during the upset period (June 15 – July 10, 2017) caused by an organic overload and nitrogen deficiency at mill C.

Table S1 Spearman correlations between Bathyarchaeia ASVs in all three mills and hydrogenotrophic and methylotrophic methanogens. These two functions were calculated for all samples using the software Faprotax (Louca et al., 2016). The table lists all significant correlations ($p<0.05$).

| ASVs | Hydrogenotrophic methanogens | Methylotrophic methanogens |
|---|---|---|
| Mill A (N=28) | | |
| Bathyarchaeia_18 | 0.59 | 0.45 |
| Bathyarchaeia_26 | 0.40 | |
| Bathyarchaeia_28 | | |
| Bathyarchaeia_33 | 0.72 | 0.62 |
| Bathyarchaeia_39 | | |
| Bathyarchaeia_40 | | 0.42 |
| Bathyarchaeia_41 | | 0.42 |
| Bathyarchaeia_47 | | 0.5 |
| Bathyarchaeia_76 | | 0.4 |
| Bathyarchaeia_81 | 0.77 | 0.41 |
| Bathyarchaeia_86 | | |
| Bathyarchaeia_88 | 0.74 | |
| Bathyarchaeia_109 | | |
| Bathyarchaeia_112 | | |
| Mill B (N=32) | | |
| Bathyarchaeia_28 | 0.44 | |
| Bathyarchaeia_32 | | |
| Bathyarchaeia_65 | | |
| Bathyarchaeia_72 | | |
| Bathyarchaeia_74 | | |
| Bathyarchaeia_76 | | 0.50 |
| Bathyarchaeia_77 | | 0.39 |
| Bathyarchaeia_86 | 0.36 | |
| Mill C (N=25) | | |
| Bathyarchaeia_17 | 0.64 | 0.52 |
| Bathyarchaeia_24 | | |
| Bathyarchaeia_36 | 0.49 | 0.63 |
| Bathyarchaeia_65 | 0.63 | 0.46 |
| Bathyarchaeia_74 | 0.63 | 0.64 |
| Bathyarchaeia_76 | 0.45 | 0.52 |
| Bathyarchaeia_77 | 0.55 | 0.73 |
| Bathyarchaeia_81 | | |
| Bathyarchaeia_86 | 0.56 | |
| Bathyarchaeia_90 | 0.46 | 0.73 |

Legend for Tables S13-S15 - List of daily operating parameters used for each mill.
Actual normalized values over time used in analyses are shown in **Excel Tables S13-S15**

#### Mill A (Table S13)

1 ORP Digester Feed
2 pH Influent
3 Temperature Influent (°C)
4 SCOD Conc. Influent (mg/l)
5 TSS Conc. Influent (mg/l)
6 pH Effluent
7 TSS Conc. Effluent (mg/l)
8 VFA-Alkalinity Ratio Effluent
9 SCOD removed (t/d)
10 OLR (kg/$m^3$/d)
11 SCOD Removal Efficiency (%)
12 $CH_4$ Conc. in Biogas (%)
13 Digester SCOD Conc. Effluent (mg/l)
14 Flow to Digesters ($m^3$/hr)
15 $SO_4$ Digester Feed
16 $SO_3$ Influent

#### Mill B (Table S14)

1 Effluent VFA-alkalinity ratio
2 Digester COD Feed Load (t/d)
3 Digester Feed SCOD Conc. (mg/l)
4 Digester 1 Effluent pH
5 Digester 1 Effluent TCOD Conc. (mg/l)
6 Digester 1 Effluent TCOD Load (t/d)
7 Digester 1 SCOD Removal Efficiency (%)
8 Digester 1 HRT (d)
9 Digester 1 Effluent TSS (mg/l)
10 Digester 2 Inf Flow (l/min)
11 Digester 2 Effluent pH
12 Digester 2 Influent SCOD Load (t/d)
13 Digester 2 Effluent SCOD Conc. (mg/l)
14 Digester 2 Effluent SCOD Load (t/d)
15 Digester 2 SCOD Removal Efficiency (%)
16 Digester 2 HRT (d)
17 Digester 2 Effluent TSS Conc. (mg/l)
18 Digester 3 Influent Flow (l/min)
19 Digester 3 Effluent pH
20 Digester 3 Inf SCOD Load (t/d)
21 Digester 3 Effluent SCOD Conc. (mg/l)
22 Digester 3 Effluent SCOD Load (t/d)
23 Digester 3 SCOD Removal Efficiency (%)
24 Digester 3 HRT (d)
25 Digester 3 Effluent TSS Conc. (mg/l)
26 Raw Biogas $H_2S$ (%)
27 Raw Biogas $CH_4$ (%)
28 $H_2S$ Load in Biogas ($m^3$/hr)
29 Influent SCOD Load (t/d)
30 Effluent SCOD Load (t/d)
31 Overall Digester SCOD Removal Efficiency (%)
32 Methane Yield daily ($m^3$/tonne TCOD)
33 Methane Yield daily ($m^3$/tonne SCOD)
34 Influent Flow to all 3 Digesters (l/hr)
35 Sulfur Load to Digesters (kg/hr)

#### Mill C (Table S15)

1 Digester Effluent COD Load (kg/day)
2 Digester TSS Conc. (mg/l)
3 Digester COD Conc. (mg/l)
4 Influent Flow to Digester ($m^3$/day)
5 Influent COD Load to Digester (kg/day)
6 Digester Influent COD Conc. (mg/l)
7 Digester Influent pH
8 Digester Influent TSS Conc. (mg/l)
9 Digester Influent Temp (°C)
10 Digester COD Removed (kg/day)
11 COD Removal Efficiency (%)
12 Anaerobic Sludge Return Line Temp (°C)
13 Methane Production ($m^3$/day)
14 Biogas Production ($m^3$/day)
15 VFA-Alkalinity Ratio
16 $H_2S$ Conc. In Biogas (%)
17 Digester pH
18 Color Conc. (CU)
